# Novel production of structurally diverse and sticky defense metabolites on wild tomatillo fruits

**DOI:** 10.1101/2025.07.16.665232

**Authors:** Lillian Nowack, Rocio Deanna, Stacey D. Smith, Craig A. Schenck

## Abstract

Plants have evolved a structurally diverse chemical repertoire to mediate various environmental interactions. Yet, little is known about the chemical complexity and metabolic pathways across many plant genera. Acylsugars are a class of specialized metabolites widely distributed across the Solanaceae that play a role in defense. Although acylsugars have been extensively characterized from leaf glandular trichomes, they have also been detected in other tissues. Nevertheless, acylsugar structures, biosynthesis, and functions outside of leaf trichomes remain unknown. Here, we performed tissue specific metabolomics across 29 *Physalis* species, an emerging model and crop genus closely related to important *Solanum* crops. Automated mass spec feature identification and library searches from surface extracts of leaf, calyx, and fruit tissues revealed metabolite diversity including putative metabolite classes of flavonoids, phenolics, and terpenoids. Acylsugar mass spec features were manually annotated, revealing at least 323 unique acylsugars - substantially expanding the known *Physalis* acylsugar diversity. Some, but not all, *Physalis* species accumulated acylsugars on the trichome-less fruit surface, and were as abundant or sometimes more abundant, in fruits than in leaves or calyces. Hierarchical clustering and phylogenetic tests indicated that species with similar acylsugar profiles do not cluster taxonomically. To determine the biochemical mechanism underlying acylsugar structural diversity, we characterized the first step of acylsugar biosynthesis, catalyzed by an acylsugar acyltransferase (ASAT). ASAT1s from three *Physalis* species displayed broad substrate preferences, which may explain the differences in acylsugar profiles. The diverse fruit-localized acylsugars across *Physalis* can inform engineering strategies for increased crop resilience.

## Introduction

Plants produce a repertoire of structurally diverse metabolites, generally divided into two classes – core and specialized (also known as secondary metabolites or natural products). Specialized metabolites largely function in abiotic and biotic stress response (Pichersky and Lewinsohn 2011; Marone et al. 2022; Wu et al. 2023) and exhibit greater structural diversity than core metabolites. These specialized metabolites are downstream of core metabolic pathways and have arisen from the co-option of duplicated core metabolic enzymes (Weng et al. 2012; Weng 2014; Fan et al. 2020). Unlike core metabolism that is conserved across plants, specialized metabolic pathways are typically lineage and tissue-specific (Pichersky and Lewinsohn 2011; Beaudoin and Facchini 2014; Schenck and Last 2020). While the ecological significance of specialized metabolites is increasingly recognized, many biosynthetic pathways and the downstream chemical diversity remain largely unexplored, particularly in non-model plants. This knowledge gap limits our ability to leverage biologically active metabolites for crop enhancement, stress resistance, and to co-opt these metabolites for medicinal and industrial purposes.

The Solanaceae family, which includes economically important species such as tomato, eggplant, potato, and petunia, is well known for its specialized chemical diversity, including terpenes, alkaloids, phenylpropanoids, and acylsugars (Fiesel et al. 2022). Acylsugars usually accumulate in glandular trichomes present on leaf and stem surfaces, and function in defense against insects, herbivores, microbial pathogens, and abiotic stressors (Weinhold and Baldwin 2011; Leckie et al. 2016; Luu et al. 2017; Feng et al. 2022). They are composed of a sugar core, usually sucrose or glucose, and acyl chains of various lengths and branching patterns that are attached at specific hydroxyl groups on the sugar core (Vendemiatti et al. 2024). Acylsugar chemical diversity has been investigated at various taxonomic levels across the Solanaceae, including the *Nicotiana* and *Solanum* genera (Schenck et al. 2022; Fiesel et al. 2024), as well as the whole family (Moghe et al. 2017). These studies have focused on trichome-localized acylsugar diversity, and a major theme is despite being assembled from simple precursors, acylsugar structural diversity is extensive. The key steps of acylsugar biosynthesis, esterification of an acyl-CoA to the sugar core, are catalyzed by a set of acylsugar acyltransferases (ASATs) that are highly expressed in the trichomes and have been biochemically characterized from a diverse array of Solanaceae species (Schilmiller et al. 2012; Fan et al. 2016, 2020; Moghe et al. 2017; Nadakuduti et al. 2017).

Despite the prevalence of acylsugars in glandular trichomes, acylsugars have been detected in other tissues, including roots in tomato and fruits in *Physalis* L. *(Zhang et al. 2016; Kerwin et al. 2024).* With roughly 90 species (Martínez et al. 2023), *Physalis* is one of the largest genera in the Solanaceae and falls within the large berry clade that also includes tomatoes, peppers, and eggplants (Särkinen et al. 2013). *Physalis* species are gaining importance as both food crops and model organisms (He et al. 2023; Lopez-Gomollon 2023; Dale et al. 2024). *Physalis* includes the crop tomatillo (*P. philadelphica* Lam.) in addition to goldenberry (*P. peruviana* L.), groundcherries (*P. grisea* (Waterf.) M.Martínez and *P. pruinosa* L.), and other species with edible fruits, medicinal properties, or grown as ornamentals (Shenstone et al. 2020; Dale et al. 2024). Mexico is the largest producer of tomatillo with an export of approximately 698,000 tons valued at 66.6 million USD in 2016 (Shenstone et al. 2020). Although a few acylsugar structures have been NMR characterized from *Physalis* fruit or calyx (Ovenden et al. 2005; Maldonado et al. 2006; Cao et al. 2015; Barrientos et al. 2022), the biological functions and distribution of fruit-localized acylsugars across *Physalis* are unknown.

Although acylsugars are best studied in vegetative tissue, their presence on the fruit surface suggests they may play novel roles in defense (i.e. protecting developing seeds). We hypothesize that fruit-surface localized acylsugars are synthesized and secreted independently from glandular trichome biosynthesis of leaf acylsugars. To gain insight into the distribution of fruit-surface localized acylsugars, we characterized acylsugar chemical diversity and biosynthesis across the *Physalis* genus. Acylsugar profiling of surface extracts revealed hundreds of acylsugar variants. Furthermore, many of the species examined accumulate acylsugars on the fruit surface, and the diversity of these compounds could not be predicted from leaf acylsugar profiles, suggesting that fruit and leaf acylsugar production may be independently regulated. The biochemical basis for acylsugar structural variation was investigated by enzymatically characterizing ASAT1 from three *Physalis* species revealing differing substrate specificities. Our findings provide valuable insights into the distribution and biosynthesis of novel fruit-surface localized acylsugars across the *Physalis* and lay the groundwork for future studies investigating their biological function.

## Results

### Metabolite profiling across Physalis reveals diverse specialized metabolites

To characterize the metabolic landscape across the *Physalis* genus, we extracted surface metabolites from three tissues, leaf, calyx, and fruit, and performed metabolomics using LC-MS. Surface extracts were prepared from 29 *Physalis* species and nine outgroup species. Some of these samples were grown from seed in growth chambers, while the majority come from extracts performed in the field in central Mexico and surrounding regions, thus not all samples have complementary leaf, calyx, and fruit samples given the nature of sample collection (Supplementary Table 1). Extracts were analyzed using UHPLC-MS primarily in negative ion mode. Mass features were identified using MSDial (Tsugawa et al. 2015) across all extracts with limited processing to attempt to capture as many unique mass spec features as possible (see methods). The number of mass spec features identified in each extract varied widely. Some extracts had as few as 500 unique mass features and as much as 8,000 mass features (Supplementary Table 2, Supplementary Figure 1). These differences likely reflect species and tissue type metabolic diversity, but also likely have to do with technical limitations and differences in extract concentrations. To assess overall metabolic diversity across tissues and species, we calculated Shannon entropy values based on the presence or absence of all mass spec features identified. *P. viscosa* had the highest Shannon entropy value, whereas *Withania riebeckii* Balf.f. had the lowest, generally consistent with the numbers of mass spec features identified and numbers of peaks observed in chromatograms (Supplementary Figure 2a and Supplementary Table 2). These data suggest that some species are enriched in metabolic complexity, at least in the extracts and tissues we analyzed. To determine if there were tissue-specific differences in metabolic complexity, we averaged leaf, calyx, and fruit Shannon entropy values across all species. Shannon entropy values for calyx and fruit were not significantly different, suggesting that their average metabolic complexity is comparable across the species analyzed (Supplementary Figure 2b, Supplementary Table 2). However, leaf metabolic complexity was significantly lower than both fruit and calyx tissue (Supplementary Figure 2b).

To determine the types of metabolites in the surface extracts, library searches were performed against the MassBank of North America (MoNA) database (Supplementary Table 1). Library matches were generally low with about 10-25% of total mass features identified using the MoNA database, with the rest remaining unidentified due to limitations in LC-MS libraries (Supplementary Figure 1). The mass spec feature library matches are not confident identities; however, they provide an overall sense of metabolite classes and metabolic diversity across tissues and species. A variety of specialized metabolite types were identified, including putative classes of flavonoids, phenolics, and terpenoids (Figure 1a, 1b). Many of these metabolites have been previously identified in various *Physalis* species: phenolics and flavonoids in *P. angulata* L.*, P. peruviana, P. hederifolia* A.Gray, and *P. patula* Mill., (Medina-Medrano et al. 2015; Huynh Nguyen et al. 2021; Kasali et al. 2021); terpenoids in *P. peruviana* and *P. angulata* (Ferreira et al. 2019; Kasali et al. 2021; Popova et al. 2022); and carotenoids and carotenoid esters in *P. peruviana* and *P. pubescens* L. (Wen et al. 2017; Kasali et al. 2021). In general, flavonoids occurred more frequently in calyx tissue, whereas phenolics were equally represented in fruit and calyx extracts (Figure 1b).

**Figure 1.**
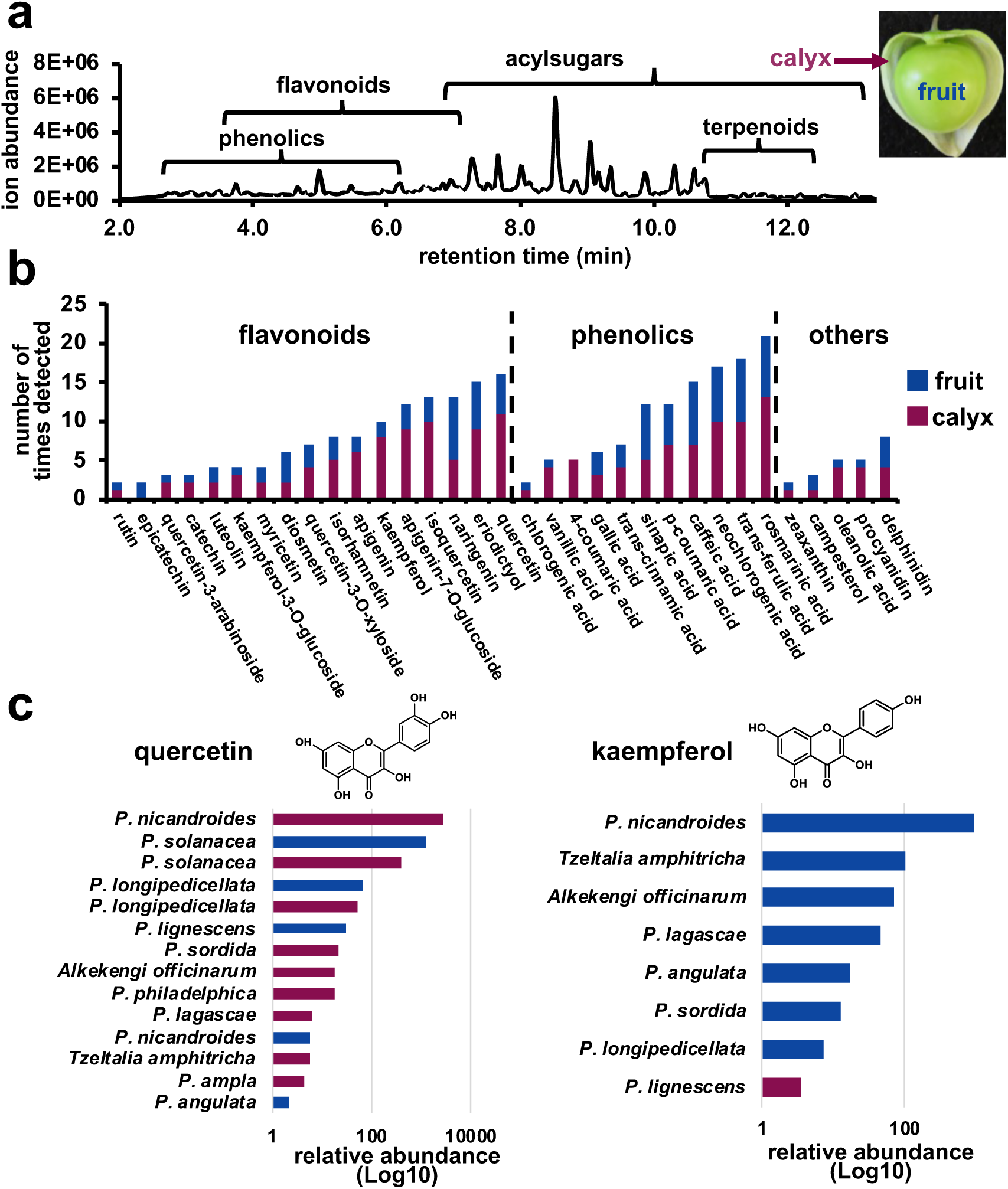
Metabolite profiling of surface extracts from *Physalis* and related species. **(a)** Representative LC-MS chromatogram showing general retention time ranges of specialized metabolite classes identified from library matches. Inset shows a *P. philadelphica* fruit with surrounding calyx tissue. **(b)** Number of times putative specialized metabolites were detected in fruit (blue) and calyx (purple) extracts. **(c)** Relative abundance of quercetin and kaempferol in a subset of extracts. Abundance represents peak areas of the MS features identified as kaempferol or quercetin and corrected for internal standard peak area and tissue dry weight. Calyx extracts in purple and fruit extracts in blue.

Next, we focused on the distribution and abundance of flavonoids given their importance in plant-environment interactions (Mierziak et al. 2014; Shomali et al. 2022), broad distribution across many plant lineages, and frequent occurrence in our dataset (Supplementary Table 2). Among the flavonoids that were detected, quercetin and kaempferol were two of the most widespread, as quercetin was present in 16 extracts, and kaempferol was present in 10 extracts from a total of 48 extracts from 27 species (Figure 1b). These metabolites have been well-studied for their antioxidant, anti-inflammatory, and antimicrobial properties (Vellosa et al. 2011; Devi et al. 2015; Tian et al. 2021; Jan et al. 2022; Periferakis et al. 2022). The relative abundance of kaempferol and quercetin was variable across species and tissues (Figure 1c). *Physalis nicandroides* Schltdl. had the highest levels of both kaempferol and quercetin whereas *P. angulata* and *P. lignescens* Waterf. had lower but still detectable levels of quercetin and kaempferol compared to other species (Figure 1c). Given that most mass spec features had no library matches, we next turned to manual annotations of a specific metabolite class to gain more information on *Physalis* chemical diversity.

### Manual annotation reveals tremendous acylsugar variation across Physalis

Although thousands of mass spec features were detected across *Physalis* species and tissues, many of the high-intensity ions were not identified using MoNa (Supplementary Figures 1 & 3a, Supplementary Table 2). Based on the mass-to-charge (*m/z*) values and fragmentation patterns, many of these features were hypothesized to be acylsugars, which are well studied within the Solanaceae but not represented in public LC-MS libraries. Acylsugar mass spec features were manually annotated in negative ion mode using the intact formate adduct mass and fragmentation patterns (Supplementary Figure 3b), yielding information about the lengths and number of acyl chains.

Manual annotation of acylsugars revealed 323 unique acylsugar types across tissues and species. (Supplementary Table 3). Acylsugars with two acylations (diacylsugars) represented 9% of the total annotated structures, whereas acylsugars with three (triacylated) and four (tetraacylated) acylations represented 63.8% and 27.2%, respectively (Supplementary Table 3, Supplementary Figure 4), suggesting that there is a biochemical bias towards the accumulation of triacylated and tetraacylated acylsugars. On a species level, accumulation patterns of di, tri and tetraacylsugars was more variable (Supplementary Figure 4). For example, *Eriolarynx lorentzii* calyx accumulated 66.7% tetraacylated acylsugars (Supplementary Figure 4). *P. aggregata* Waterf., on the other hand, accumulated two diacylated, seventeen triacylated and four tetraacylated, consistent with the overall acylation trends observed (Supplementary Figure 4). In contrast, many of the 323 manually annotated acylsugars had identical annotations but differing retention times, indicating the presence of isomers differing in either acyl chain positioning on different hydroxyl groups or acyl chain branching pattern. Two annotations were particularly rich in isomers, S3(21):4,5,12 and S3(22):5,5,12, where S indicates sugar core (sucrose in this case) with three acylations: totaling 20 and 21 carbons, respectively and with the individual acyl chain lengths of 4, 4, 12 and 4, 5, and 12 carbons each. S3(21):4,5,12 had 14 distinct retention times and S3(22):5,5,12 had 15 (Supplementary Table 3).

To complement the negative mode annotations, positive mode analysis was performed for a subset of samples (Supplementary Figure 3c, Supplementary Table 3). Positive mode annotations generally confirmed negative mode annotations and provided evidence for locations of acyl chains on the pyranose (six-membered) or furanose (five-membered) ring. Positive mode annotations revealed that the long acyl chain, usually 10 or 12 carbons long, was usually attached to the pyranose ring of the sugar core (Supplementary Table 3, Supplementary Figure 3c). The acyl chain attachment was consistent with previously elucidated NMR structures of *Physalis* acylsugars (Ovenden et al. 2005; Maldonado et al. 2006). However, the short and medium chains were more variable on whether they were attached to the pyranose or furanose ring of the sugar core, possibly suggesting enzymatic flexibility for other steps in the pathway.

Although fragmentation patterns yielded ions lacking acyl chains consistent with a disaccharide sugar core in many *Physalis* extracts, the identity of the sugar core was not confirmed from LC-MS fragmentation patterns alone. Based on previously characterized NMR structures from *Physalis* species, we hypothesized that the disaccharide core present across most *Physalis* is sucrose (Ovenden et al. 2005; Maldonado et al. 2006; Cao et al. 2015; Zhang et al. 2016; Barrientos et al. 2022). To confirm the sugar core type, acylsugars were extracted from a subset of species (*P. philadelphica, P. hederifolia, P. heterophylla* Nees), the acyl chains removed, then the remaining sugar core was derivatized and injected into GC-MS. *Solanum pennellii* Correll was used as a positive control, as it has been shown to produce both acylglucoses and acylsucroses (Lybrand et al. 2020). Our analysis confirmed the presence of both glucose and sucrose cores in *S. pennellii* Correll acylsugars (Supplementary Figure 5). Only sucrose was detected in *P. philadelphica,* consistent with LC-MS acylsugar annotations (Supplementary Table 3). Based on LC-MS annotations, *P. hederifolia, P. heterophylla* Nees produce primarily acylsucroses (Supplementary Figure 5). GC-MS analysis of *P. hederifolia, P. heterophylla* Nees cores reveals predominant sucrose and minor glucose peaks (Supplementary Figure 5). While sucrose is the predominant core for acylsugars in *Physalis,* specific lineages and tissues may produce acylglucoses, providing more variation to the overarching chemical diversity.

*Physalis* species produced acylsugar chains with as few as 2 carbons and up to 12, with most acylsugar structures consisting of at least one medium-chain acyl chain (C8-C12), and several short chains (C2-C6) (Supplementary Table 3, Supplementary Figure 6). LC-MS fragmentation pattern does not provide information about branching patterns of acyl chains or locations of potential double bonds. To provide additional structural information about acylsugar acyl chains, GC-MS analysis was performed on a subset of species with abundant acylsugars. GC-MS revealed the presence of several branched chains such as isobutyryl (iC4), isovaleryl (iC5), and isohexanoyl (iC6) (Supplementary Figure 7), which arise from branched-chain amino acid precursors such as valine, isoleucine, and leucine (Slocombe et al. 2008). Detectable levels of desaturated C5 were also present in these species, but generally in low amounts. The longer acyl chains (C8-C12) were typically straight and unsaturated (Supplementary Figure 7), in agreement with the previously elucidated NMR structures (Ovenden et al. 2005; Maldonado et al. 2006; Cao et al. 2015; Barrientos et al. 2022). Various complementary analytical approaches confirm the presence of a large degree of acylsugar structural variation across the *Physalis*.

### Tissue-specific variation in acylsugar accumulation

Although a handful of acylsugar structures have been NMR characterized from *Physalis* fruit (Maldonado et al. 2006; Zhang et al. 2016), we wanted to understand how widespread the fruit-localized acylsugar trait is across *Physalis*, as it appears to be a unique location for acylsugar accumulation. To evaluate tissue-specific acylsugar differences, we compared the abundance and number of acylsugars in calyx and fruit tissues across a subset of *Physalis* species (Figure 2).

**Figure 2.**
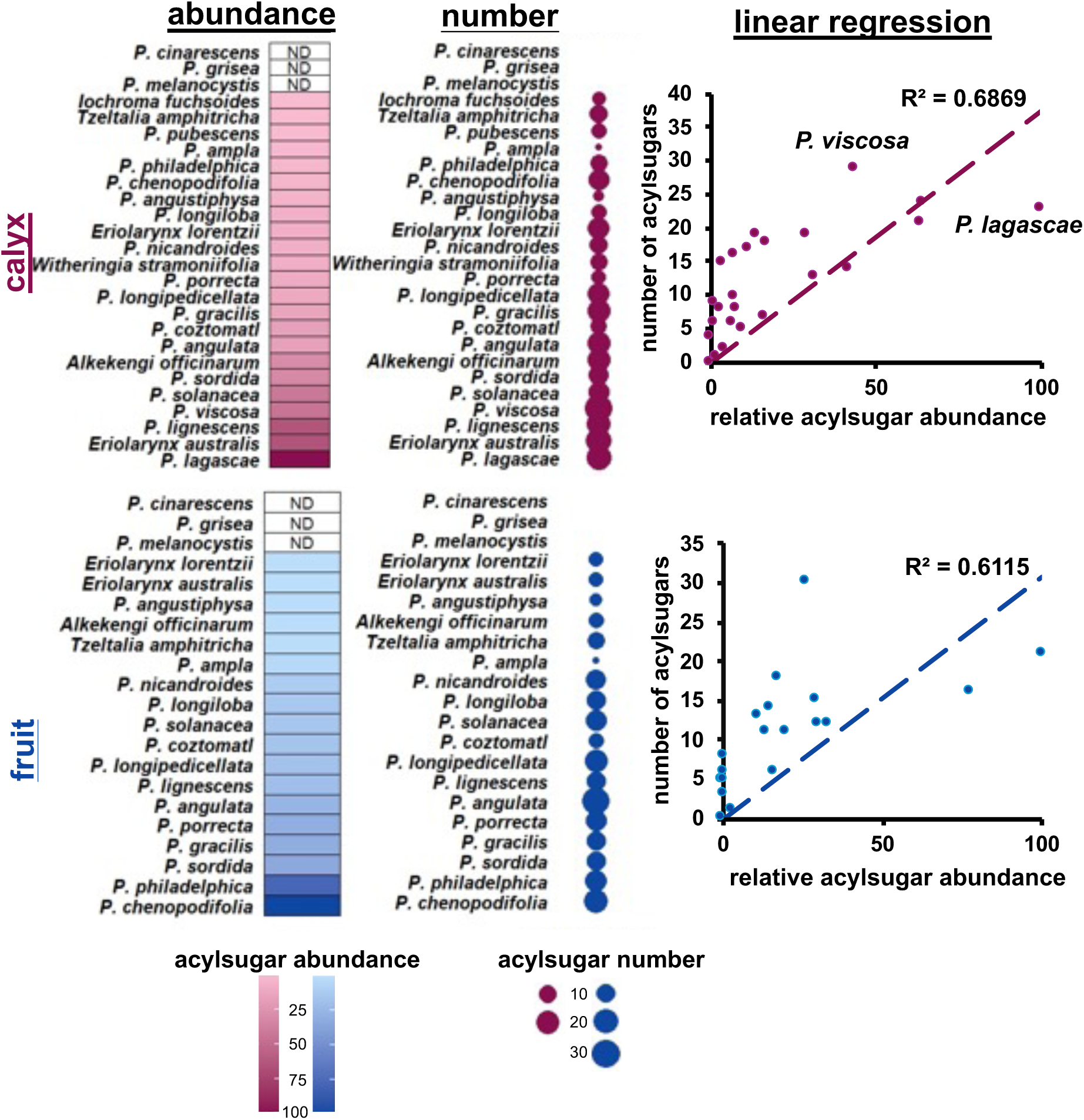
Correlation of number and abundance of *Physalis* fruit and calyx acylsugars. Acylsugars were quantified using the total peak area of all annotated acylsugars corrected for internal standard peak area and tissue dry weight. In a tissue specific manner, the most abundant extract was set to 100% and all other extracts were normalized to this value. N.D., below detection limit. Size of circle indicates umber of acylsugar types in calyx and fruit. Lack of circle indicates no detectable acylsugars from that tissue. Linear regression of the relationship between relative abundance and number of acylsugars, with R^2^ values for line of best fit shown.

Fruit localized acylsugars were detected from 22 species, whereas seven species had no detectable fruit localized acylsugars (Supplementary Figure 8). While many species produced abundant fruit and calyx acylsugars, acylsugars were not detectable from leaf extracts of most *Physalis* species profiled, except for *P. pruinosa, P. heterophylla,* and *P. hederifolia* (Supplementary Table 3, Supplementary Figures 8 & 9). This is supported by glandular trichome counts, as most species had few to no glandular trichomes on the leaf surfaces used for acylsugar extractions (Supplementary Figure 10a). Both acylsugar abundance and number varied widely across species and tissue types (Figure 2). *Physalis ampla* produced only one annotatable acylsugar in fruit tissue; however, *P. angulata* produced 30 (Figure 2). Generally, acylsugar profiles were similar in calyx and fruit tissues, with fruit producing a larger number of annotatable acylsugars, possibly due to fruit extracts having more abundant acylsugars (Supplementary Figure 11). To test if calyx acylsugar abundance is correlated with fruit acylsugar abundance we performed a regression analysis between fruit and calyx acylsugar traits. We find a lack of correlation between these two traits, suggesting independent control over fruit and calyx localized acylsugar accumulation (Supplementary Figure 12a). A parallel analysis incorporating phylogenetic effects also failed to recover a relationship between fruit and calyx acylsugar abundance across species (Supplementary Figure 12b). While many species had abundant acylsugars on the fruit surface, it remains unclear whether the acylsugars are being biosynthesized on the fruit surface, the calyx, or both.

To assess the relationship between the number and abundance of acylsugars, linear regression was performed with the normalized abundance values and number of acylsugars for all samples in a tissue-specific manner. The coefficients of determination (R^2^) values for calyx and fruit were 0.69 and 0.6, respectively (Figure 2), indicating a moderately positive correlation between the abundance and number of acylsugars. Some species deviated from this trend. For example, *P. philadelphica* fruit displayed a high abundance but less in regard to number, while others, such as *P. viscosa* calyx, showed the inverse. These patterns suggest that while abundance is influenced by the number of acylsugars produced, there are other contributing factors.

### Acylsugar structural diversity in relation to the phylogeny

Next, we wanted to examine the degree to which acylsugar production relates to phylogenetic relationships among the species. To determine the similarity of acylsugar profiles across species and tissues, we first carried out hierarchical clustering analysis (HCA) and principal coordinate analysis (PCoA) based on the absence and presence of each of the 323 manually annotated acylsugars. Out of the 323 annotated acylsugars, 237 (∼73%) were detected in only one species, highlighting the predominance of species-specific acylsugars (Figure 3). Furthermore, only 40 were shared across three or more species. This indicates limited overlap in acylsugar production across the genus, despite their close evolutionary relationship. Moreover, the patterns of clustering recovered from HCA (Figure 3b) were only loosely tied to the taxonomic groupings. In particular, some of the outgroup taxa (*Eriolarynx* (Hunz.) Hunz., *Iochroma* Benth.) did cluster separately from the rest of the species (Figure 3b), but others appear very similar to some *Physalis* species. To further explore acylsugar variation, PCoA was conducted using the same binary acylsugar matrix. Similar to HCA, the PCoA highlighted distinct clusters of species that have similar acylsugar profiles (Supplementary Figure 13). Here again, the *Iochroma* and *Eriolarynx* samples appear very distinct from the rest, while *Witheringia* and *Tzeltalia* grouped with *Physalis* species.

**Figure 3.**
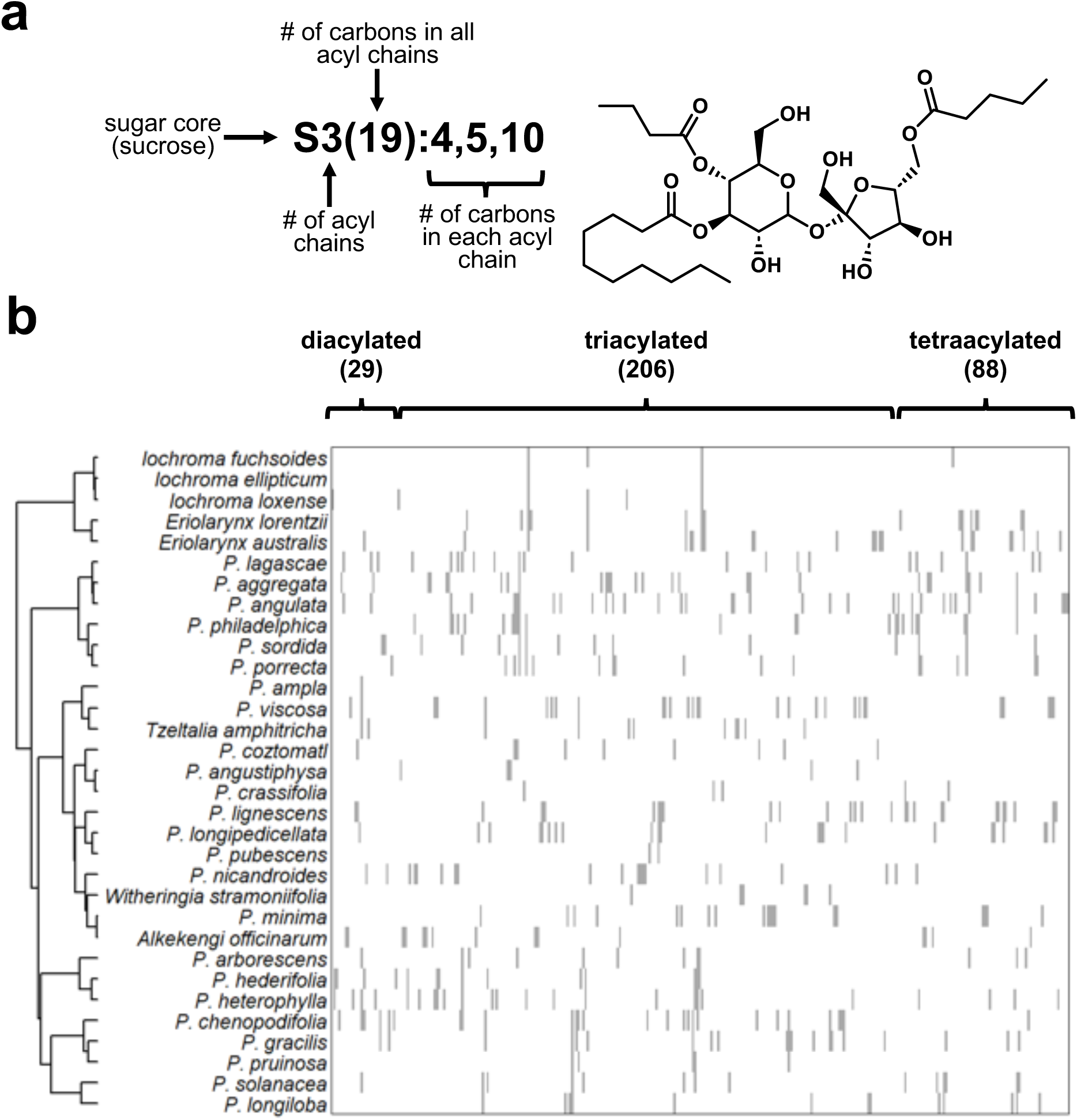
Hierarchical clustering of acylsugar structural variation across all species. **(a)** Acylsugar nomenclature used in this and other studies with a representative triacylated structure shown. **(b)** Hierarchical clustering analysis (HCA) of presence/absence of each annotated acylsugar. Each row represents a species, while each column represents a single acylsugar annotation in order of number of acylations, then increasing size in total number of carbons in the acyl chains. Number of diacylated, triacylated, and tetraacylated acylsugars identified are shown in parenthesis. Grey indicates presence of an acylsugar, white indicates absence. Acylsugar presence for each species represents the sum of all acylsugar types present across all tissues.

To more rigorously assess the relationship between chemical profiles and the phylogeny, we computed two measures of phylogenetic structure in the acylsugar data and tested for significant signal. First, we calculated Blomberg’s (Blomberg et al. 2003) for acylsugar abundance in fruits and calyces using a sample of 100 Bayesian trees from the phylogenetic analysis of Deanna et al. 2019. Values of K ranged from 0.16 to 0.64 across trees and traits, and none were significantly different from zero (Supplementary Table 4), suggesting no detectable phylogenetic signal. We found a similar lack of signal in the type of acylation. We scored each of the species as present or absent for each class of acylsugars (di, tri, tetra) in any tissue and used Fritz and Purvis’s D statistic (Fritz and Purvis 2010) to test for signal in these binary variables. Again, using the sample of 100 trees, these three traits presented values of D closer to 1, which is the expected value from no phylogenetic structure (Supplementary Table 5). It is worth noting, however, that many of the sampled species are not included in the phylogeny (e.g., only 18 species are present in the phylogeny and have calyx acylsugar abundance data), and generally the power to detect signal is low with fewer than 20 species (Blomberg et al. 2003).

### Identification and biochemical characterization of Physalis ASAT1s

To investigate the biochemical basis of the chemical diversity, we identified and characterized three ASAT1s from different *Physalis* species with contrasting acylsugar profiles: *P. pruinosa, P. philadelphica,* and *P. coztomatl* Dunal. ASAT1 candidate genes were identified using biochemically characterized *Solanum lycopersicum* L. ASATs to BLAST the *P. pruinosa* genome. BLAST hits were filtered based on the presence of two conserved domains present in BAHD acyltransferases, HxxxD and DFGWG motifs. A phylogenetic analysis was performed with *P. pruinosa* candidates together with biochemically characterized ASATs from Solanaceae species (Supplementary Figure 14). One *P. pruinosa* candidate grouped within a clade containing previously characterized ASAT1s called ancestral ASAT1 (AncASAT1) as they are present in most Solanaceae lineages but absent in *Solanum* (Supplementary Figure 14). Primers designed for *P. pruinosa* ASAT1 were used to amplify and clone *P. pruinosa* ASAT1 as well as homologous ASAT1 sequences from *P. philadelphica* and *P. coztomatl. Physalis* ASAT1 enzymes were cloned, recombinantly expressed, and purified for biochemical analyses. Substrate specificity of *Physalis* ASAT1s was assessed *in vitro* using commercially available acyl-CoAs of varying lengths together with sucrose. *In vitro* assays revealed slight differences in substrate specificity. While all three ASAT1s preferred C12 CoA, *P. philadelphica* had a narrower substrate specificity range than the other two *Physalis* ASAT1s (Figure 4a). The preference for the C12 product is consistent with the widespread presence of long-chain acylsugars (Supplementary Figure 6), as well as with the presence of a long chain in most NMR elucidated structures (Ovenden et al. 2005; Maldonado et al. 2006; Cao et al. 2015; Barrientos et al. 2022).

**Figure 4.**
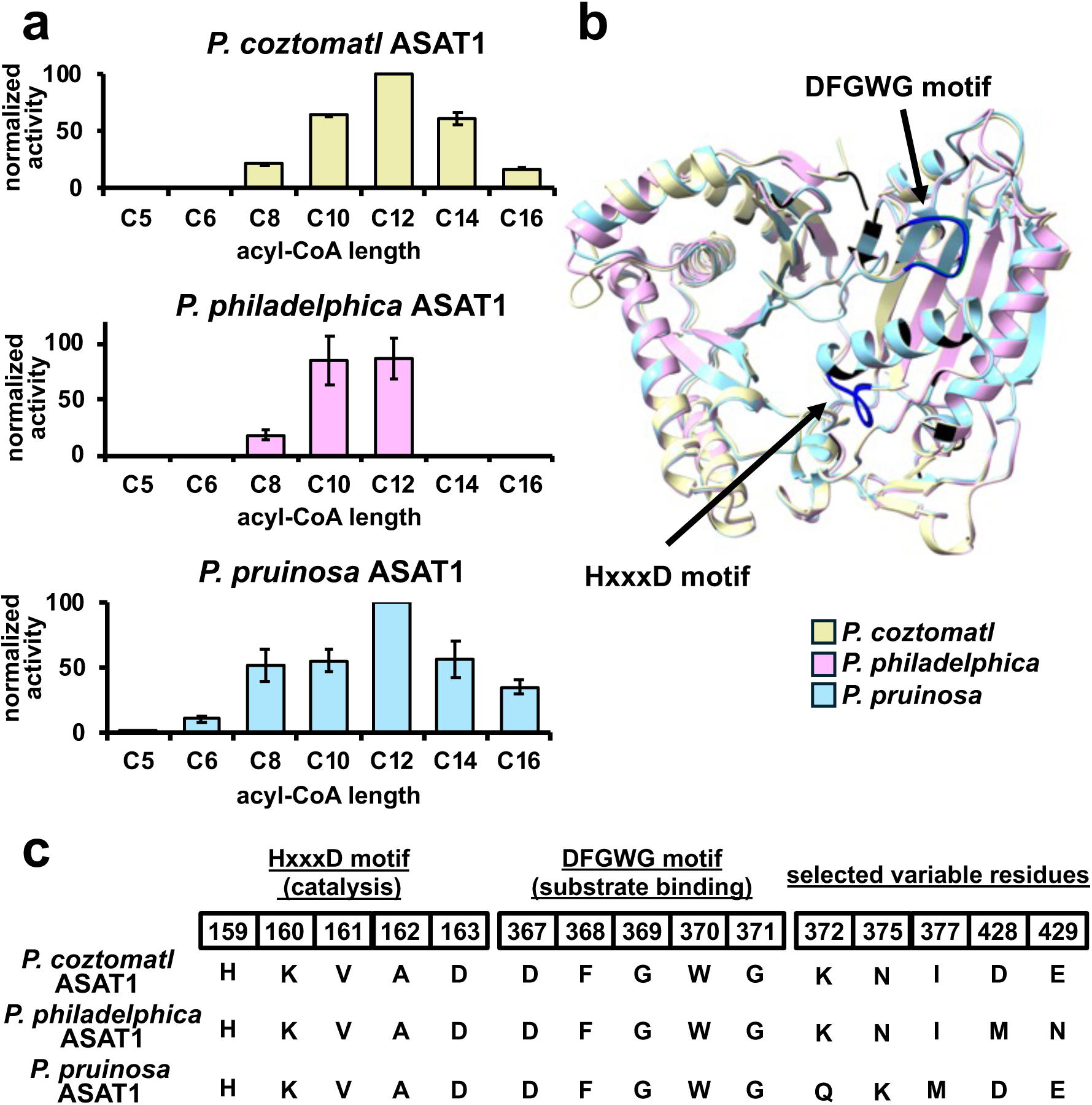
*Physalis* ASAT1 *in vitro* biochemical and structural characterization. **(a)** Substrate promiscuity of *Physalis* ASAT1s. Enzyme activity was tested with purified recombinant enzymes with varying lengths of acyl-CoA (250µM) and sucrose (2mM) and monitored for production of monoacylated products using LC-MS. Values are means ± standard deviation of two biological replicates. **(b)** Structural model overlays of ASAT1s, with amino acid changes in black, and catalytic motifs in dark blue. **(c)** Amino acid alignment of conserved and variable regions of ASAT1s. Amino acid numbering based on *P. pruinosa* ASAT1.

To explore the biochemical basis for changes in substrate specificity, AlphaFold structural models were generated for *Physalis* ASAT1s. The ASAT1s showed sequence variation, with 12 amino acid changes between *P. philadelphica* ASAT1 and *P. coztomatl* ASAT1, 28 between *P. pruinosa* ASAT1 and *P. philadelphica* ASAT1, and 30 between *P. pruinosa* ASAT1 and *P. coztomatl* ASAT (Supplementary Figure 15). Structural modeling revealed slight changes in the tertiary structure of these enzymes, primarily at the N terminus, and slight changes in the positioning of alpha-helices (Figure 4b). Several of these changes are near conserved HXXXD and DFGWG binding motifs, necessary for catalysis and structural stability of BAHD family acyltransferases (D’Auria 2006) and may be responsible for the substrate specificity differences observed across *Physalis* ASAT1s (Figure 4c).

## Discussion

### Metabolic diversity in Physalis species

Metabolic profiling across the *Physalis* genus revealed a wide array of specialized metabolites, including acylsugars and putative flavonoids, phenolic acids, and terpenoids (Figures 1, 2 & 3), highlighting the extensive metabolic diversity across the genus. These specialized metabolite classes play important roles in plant defense and other ecological interactions – such as attracting pollinators, deterring herbivores, protecting from UV damage, and reducing water loss (Erb and Kliebenstein 2020). The *Physalis* genus is well known for a unique morphological trait – an inflated calyx surrounding the fruit – that has been hypothesized to serve a number of ecological and physiological functions, including stabilizing the microclimate during fruit development (Li et al. 2019), aiding in dispersal (Wilf et al. 2017), and protecting the berry (de Souza et al. 2022). The presence of fruit-surface localized acylsugars and morphological innovations suggests the presence of a multilayered defense response in *Physalis* species.

### Acylsugar diversity and biological activity implications

Acylsugars in *Physalis* displayed extensive structural diversity. Over 300 distinct acylsugars were annotated, of which only ∼80 were present in more than one species, showing that the majority of acylsugar structures are species-specific (Supplementary Table 3, Figure 3). Because of technical limitations in detection limits and separation of metabolites, the number of acylsugars across *Physalis* is much greater than what can be annotated from LC-MS experiments, possibly more than 600 distinct structures (Yuan et al. 2025). It remains unclear as to why so many distinct acylsugar types accumulate across *Physalis* and the Solanaceae.

Producing a repertoire of acylsugar types may provide a broad layer of defense that is challenging to evolve an avoidance mechanism. Acylsugars have demonstrated roles in biotic and abiotic stress response (Vendemiatti et al. 2024a) and some acylsugars with long chains have been shown to have a greater effect on certain insects (Puterka et al. 2003). Thus, different acylsugar types may have specific functions (e.g., some may be more useful in abiotic stress tolerance and others for biotic stresses).

### ASAT promiscuity enables acylsugar structural variation

Despite the evolutionary relationships among the sampled species, there are many differences in the overall acylsugar profile, abundance, and tissue-specificity. *Physalis* ASAT1s grouped within a clade containing ancestral ASAT1s from most Solanaceae lineages excluding *Solanum* ASAT1s (Supplementary Figure 14). Although Ancestral ASAT1s and *Solanum* ASAT1s acylate at different hydroxyl groups they both have broad substrate specificity profiles for the acyl-CoA substrate (Fan et al. 2016; Moghe et al. 2017; Schenck et al. 2022). Acylsugar structural diversity reflects the substrate promiscuity of ASATs, but also the availability of acyl-CoA substrates at the location of biosynthesis among other factors (Fan et al. 2017; Schenck et al. 2022). This is highlighted especially by the substrate specificity and slight structural changes in the first step of the acylsugar biosynthetic pathway, catalyzed by *Physalis* ASAT1s (Figure 4). The broad substrate profile observed in *Physalis* ASAT1s (Figure 4) is a likely driving force in the overall numbers of acylsugars produced. Amino acid alignments and structural modeling highlight some amino acids that could play a role in the range of substrate preferences observed across *Physalis* ASAT1s (Figure 4, Supplementary Figure 15), which could be investigated using site directed mutagenesis. Given the broad Acyl-CoA substrate profiles typical of ASATs, it could be possible to engineer ASATs for greater specificity to generate acylsugar types with increased activities.

### Acylsugars vary independently of the phylogeny

Although we are only just beginning to understand the diversity of acylsugar production in *Physalis*, our results suggest that this diversity is not strongly predicted by evolutionary history. Except for species belonging to the Iochrominae subtribe (*Eriolarynx* and Iochroma), our clustering analyses based on chemical profiles group species together that are not closely related on the phylogeny (Figure 3, Supplementary Tables 4 & 5, Supplementary Figures 12 & 13,). This is partly because most species accumulate distinct, species-specific acylsugar types, with only a handful of compounds shared across species (Figure 3). In addition, when we directly estimate phylogenetic signal in acylsugar abundance and type, we find no significant phylogenetic signal (Supplementary Tables 4 & 5). Together, these results indicate that acylsugar profiles are not a good indicator of species relatedness. Nevertheless, thirteen of our study species are not present in the most recent phylogeny for the group (Deanna et al. 2019). A more resolved and comprehensive *Physalis* phylogeny together with additional acylsugar profiling from more *Physalis* would provide a clearer picture of evolutionary trends in acylsugar production and allow us to more confidently estimate the tempo and mode of changes in acylsugar traits such as acyl chain lengths, core types, and tissue specific accumulation.

### Widespread fruit-localized acylsugars in Physalis

Canonically, acylsugars are synthesized in the glandular trichomes on leaf and stem surfaces (Vendemiatti et al. 2024b). In most *Physalis* species screened, there were no detectable acylsugars in leaves (Supplementary Figure 9), supported by few to no glandular trichomes on the leaf surface (Supplementary Figure 10a). However, we identified many species producing fruit-localized acylsugars, a unique observation especially considering the lack of glandular trichomes on the fruit surface (Supplementary Figure 10b). Fruit-localized acylsugars were identified in 22 of the 39 species profiled (Supplementary Figure 8), highlighting how widespread this trait is within the genus, and indicating this may be evolutionarily advantageous. In many cases, acylsugars were more abundant on the fruit-surface as compared to the other tissues screened, suggesting the novel localization serves an important function. Fruit-surface localized acylsugars are sticky, like trichome-localized acylsugars, and thus could serve as a trap for small insects but additionally could serve to protect the fruit from desiccation and the developing seeds from biotic stressors. Moreover, acylsugar production in fruits appears to evolve independently from production in these other tissues (Supplementary Figure 12), suggesting tissue-specific transcriptional mechanisms that could be leverage in the future for targeted crop improvements.

Despite widespread distribution of fruit localized acylsugars across the *Physalis* it is unknown how they are deposited on fruit surface. Knowing the biosynthetic pathway will provide tools to investigate their expression in different tissues and locations. It is possible that ASATs are expressed in modified trichomes or secretory cells on the fruit surface, analogous to how volatile compounds are secreted from citrus fruit peels (Voo et al. 2012; Wang et al. 2024). Further work is required to understand the biosynthesis and deposition of fruit-surface localized acylsugars.

This knowledge can be used to remove the stickiness from crop fruits if it is not required for protection, or this trait could be engineered in other berry crops, like tomato, for enhanced protection.

## Materials and methods

### Field collections and plant growth

Field extractions were performed during collection trips in central and Sourthern Mexico and central Argentina, except *W. riebeckii* which was collected at the Kaisaniemi Botanic Garden (Helsinki, Finland). Plants were identified by R. Deanna, P. Zamora-Tavares, A. Herrera, O. Vargas-Ponce, and M Martínez, and voucher specimens were stored at the CORD, H, IBUG, and QMEX herbaria.

When the extractions were not performed in the field, we cultivated the plants in the greenhouse. *Physalis* seeds were obtained from the US Department of Agriculture Germplasm Resources Information Network (GRIN) or from previous collection trips. Seeds were treated with 10% sodium phosphate tribasic and 50% bleach (v/v) solution for 5 minutes with constant rocking.

Seeds were rinsed five times with distilled water, placed on Whatman filter paper, and germinated at 28°C in the dark. Seedlings were transplanted into soil and grown at 27°C during the day and 22°C during the night under a 18/6 day/night cycle with a light intensity of ∼9,000 lux. Plants were fertilized once per week with MiracleGrow.

### Surface metabolite extraction, quantification, and detection

Metabolites used for LC-MS analysis were extracted from young leaf tissue or mature fruit and calyx using 3:3:2 isopropanol:acetonitrile:water with 0.1% formic acid (Vendemiatti et al. 2024a). Tissue was submerged in 1mL of extraction solvent and gently agitated for 3-5 minutes. The extraction solvent was dried to completion using a centrivap (ThermoScientific) then resuspended in 100µL of extraction solvent with a final concentration of 3µM telmisartan internal standard. Extracts were placed in a 2 mL glass vial with an insert. 4uL of the extracts were injected into a Bruker Compact LC-MS qTOF with an Ascentis Express 90 Å C18 column for reverse-phase separation with the column temperature set at 30°C. The starting concentration was 95% solvent A (10mM ammonium formate, adjusted to pH 2.8 with formic acid) and 5% solvent B (100% acetonitrile) with flow rate set to 0.3 mL/min. Metabolites were separated on a C18 reverse-phase column using a 20-minute LC gradient (Supplementary figure 16) using an UltiMate 3000 system operating in positive and negative ion modes.

Putative acylsugars were identified that have a retention time between 6.0 and 15.5 minutes, *m/z* between 639 and 835, and adducts of either [M-H]^-^ or [M+FA-H]^-^. *m/z* values were corrected using the sodium formate injection during each LC-MS run. Full scans were collected from 15 to 1500 *m/z*. Mass spec features were identified in MS-DIAL as [M-H]^-^, [M-2H]^2-^, or [M+FA-H]^-^ adducts with a minimum peak heigh of 500 amplitude, and annotated by searching against MassBank of North America database with a MS1 mass tolerance of 0.01 Da and without MS2 spectra. Manual annotation was performed on these putative acylsugars, and those that were annotated were used for quantification. Acylsugars were quantified by summing all peak areas for identified acylsugars from LC-MS analysis performed in MS-DIAL. The total acylsugar peak area was corrected for the peak area of internal standard (telmisartan) and by tissue dry weight in grams. The resulting values were normalized to the most abundant extract for each tissue type.

### LC-MS positive and negative mode annotation of acylsugars

Acylsugar structures were inferred from *m/z* of intact adducts (either formate in negative mode or ammonium in positive mode) and fragmentation patterns in negative mode and positive mode (Supplementary Figure 3) as previously described (Lybrand et al. 2020; Fiesel et al. 2024).

Acylsugars were considered the same if the retention time (RT) was within 0.05 minutes and mass-to-charge ratio (*m/z*) was within 0.1 *m/z*. Positive ion mode fragmentation was used to infer acyl chain positioning on the glucose or fructose monomer of the sucrose core, due to the typical cleavage of the glycosidic bond of the sucrose before the ester bond of the acyl chains (Lybrand et al. 2020). Acyl chains were annotated as described in Supplementary Figure 3. Acylsugars with similar *m/z* yet differing retention times were considered to be different acylsugars, as this indicates the presence of isomers. Those with similar retention times, exact masses, and fragmentation patterns were considered to be the same acylsugar.

### Shannon entropy

Shannon entropy was calculated based on the presence or absence of all detected metabolites. Each detected metabolite feature was defined by its unique combination of its *m/z*, retention time, and adduct type. To account for instrument variation and to align features across samples, *m/z* and RT values were rounded to the nearest tenth prior to feature identification. Features sharing identical rounded *m/z*, RT and adduct type values were treated as the same metabolite. Metabolite presence was determined for each sample, regardless of intensity. Shannon entropy was then computed via R using the formula:

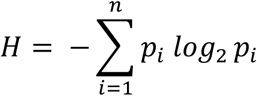

where *p_i_* is the relative frequency of each mass spec feature (in this case all equal due to just using absence and presence), and *n* is the total number of mass spec features in the sample. Entropy values were then sorted by species and tissue type, calculating mean entropy across samples. All data processing and entropy calculations were performed in RStudio using the readxl, dplyr, tidyr, ggplot2, multicompView (Wickham 2009).

### PCoA and HCA

Acylsugar annotation data were compiled into a binary absence (0) and presence (1) matrix, with rows representing species and columns representing unique annotations. Acylsugars with the same annotation but differing retention times were treated as unique annotations. Pairwise species dissimilarities were computed using Jaccard distance. Principal Coordinates Analysis was performed on the dissimilarity matrix, and the first two principal coordinate axes (PCoA1 and PCoA2) were visualized in a scatter plot. Each point represents a species, notated by the specific color and shape. These analyses were carried out using the readxl, dplyr, vegan, ape, and ggplot2 packages in R (Wickham 2009; Paradis and Schliep 2019).

Hierarchical clustering analysis (HCA) was performed with the same binary absence and presence matrix as was used for Principal Coordinate Analysis (PCoA). Pairwise dissimilarity between species was computed using Jaccard distance, then the Jaccard distance matrix was projected into principal coordinate space. A new distance matrix was made from the top five PCoA axes, then hierarchical clustering performed using Ward’s minimum variance method. The dendrogram was used to order species in a heatmap showing the absence (white) and presence (grey) of annotations. Annotations retained their original order – sorted first by number of acylations then by *m/z.* These analyses were carried out using the readxl, dplyr, vegan, ComplexHeatmap, and circlize packages in R (Gu et al. 2014, 2016; Gu 2022).

### Measures of phylogenetic signal

We estimated two measures of phylogenetic signal, namely Blomberg’s K (Blomberg et al. 2003) for the continuously varying acylsugar abundance, and Fritz and Purvis’s D (FPD, Fritz and Purvis 2010) for binary acylation type (e.g. presence/absence of diacylation). We used a sample of 100 Bayesian time trees for Physalideae from Deanna et al. (2019). We pruned the trees to include only the species with trait and the trait data to include only taxa in the trees. For K, we used 1000 simulations to test for phylogenetic signal (K significantly greater than zero) and for D, we carried out 1000 permutations to estimate the probabilities of observing the data assuming no effect of phylogeny and of observing the data with evolution along the phylogeny following the Brownian motion process. We report mean, maximum and minimum values for K and D for each trait across the 100 trees (Supplementary Tables 4 & 5) as well as the mean, maximum and minimum P-values (for K) and mean, maximum and minimum of P0 and P1 (for D). These analyses were carried out using the phytools 2.1-1(Revell 2024) and caper 1.0.3 packages for R (Orme 2023).

### GC-MS acyl chain analysis

Acylsugars were extracted as previously described using 3:3:2 isopropanol:acetonitrile:water with 0.1% formic acid in bulk. Extracts were dried to completeness in a centrivap. To cleave acyl chains from acylsugars and convert them into ethyl esters, 300μL of 21% (v/v) sodium ethoxide was added to the dried bulk acylsugar extract. Samples were incubated at room temperature for 30 minutes with vortexing every 5 minutes. To isolate ethyl esters, 400µL of hexane was added followed by vortexing and then 500µL saturated sodium chloride was added to make a phase separation. The hexane layer was then transferred to a clean tube and additional sodium chloride was added. The hexane layer was moved to a fresh tube followed by sodium chloride. The final hexane layer was then moved to a 2mL glass vial with a glass insert (Schenck et al. 2022). One microliter of sample was injected with a 10:1 split ratio onto a 60m DB-5ms column (Agilent Technologies). The initial oven temperature was 120°C, with an oven ramp rate of 6°C per minute until 300°C and a hold time of 8 min, and the inlet valve temperature was maintained at 280°C. The GC-MS was operated in full scan mode from a mass-to-charge ratio (*m/z*) of 50–650. Commercially available ethyl ester standards (Sigma) were used to aid in peak identification. Peaks absent in the even chain fatty acid ethyl ester standards (such as iC4, aiC5, iC5, and desaturated C5) were identified based on spectral matches to the NIST database.

### Sugar core analysis

Bulk extracts from a subset of *Physalis* species were dried to completion in a centrivap, then resuspended with 0.25mL H_2_O and 1mL chloroform. Aqueous and organic phases were allowed to separate, then the organic phase was transferred to a separate tube and dried at room temperature overnight. Once evaporated, the dried sample was resuspended in 100µL of isopropanol. For saponification, 20µL of sample was added to a tube with 200µL methanol and 200µL 3N ammonium hydroxide. Sample was vortexed to mix then incubated at room temperature for 48 hours. Tubes were then dried to completion overnight in a centrivap. The dried extract was resuspended in 50µL of a pyridine solution containing 15 mg/mL of methoxyamine HCl and incubated for 1 hour at 50°C, then 50 mL of *N,O*-Bis(trimethylsilyl)trifluoroacetamide + 1% trimethylchlorosilane (BSTFA + 1% TMCS; Supelco) was added and incubated for 1 h at 50°C (Schenck et al. 2020; Gibson et al. 2025).

Derivatized samples were injected into an Agilent 5977C GC-MS with authentic sugar standards (glucose, sucrose and *myo*-inositol). One microliter of sample was injected with a 10:1 split ratio onto a 60m DB-5ms column (Agilent Technologies). The initial oven temperature was 120°C, with an oven ramp rate of 6°C per minute until 300°C and a hold time of 8 min, and the inlet valve temperature was maintained at 280°C. The GC-MS was operated in full scan mode from a mass-to-charge ratio (*m/z*) of 50–650.

### ASAT identification and purification

ASATs in *Physalis* were identified using *P. pruinosa* genome. ASAT1-4 from *Solanum lycopersicum* were used to BLAST the *P. pruinosa* genome (He et al. 2023). 157 candidates were identified, which were filtered based on size, as ASATs are typically around 400-460 amino acids, as well as the presence of the HXXXD and DFGWG domains required for catalysis and substrate binding, respectively (Xu et al. 2023). The 40 remaining candidates were grouped with biochemically characterized ASATs from other Solanaceae species (Fan et al. 2016; Moghe et al., 2017; Schenck et al. 2022) in a phylogenetic analysis using the Jukes-Cantor Distance model and neighbor-joining tree build method with 100 bootstrap replicates. A single candidate emerged based on the phylogenetic positioning with other ancestral ASAT1s (Supplementary Figure 14), as well as its high bit-score and E-value.

Forward and reverse primers were designed based on *P. pruinosa* ASAT1, and were used to amplify ASAT1 from *P. pruinosa, P. philadelphica,* and *P. coztomatl* genomic DNA, as ASATs typically do not have introns. Polymerase chain reaction (PCR) was performed using ThermoScientific Phusion DNA polymerase with a primer annealing temperature of 56°C, extension temperature of 72°C, and a total of 34 cycles. Primer pair used for gene amplification included the forward primer 5’-GCTAGCATGACTGGTGGACAGCAAATGGCTGCCTCAGCTCTA-3’ and reverse primer 5’-GCCGGATCTCAGTGGTGGTGGTGGTGGTGCTTATTCATCAACAAGTTTGAAT-3’.

Amplified genes were purified by agarose gel electrophoresis and gel extraction using the Qiagen QIAquick gel extraction kit. ASAT candidate genes were cloned into pET28b expression vector through Gibson assembly according to manufacturer’s instructions. Cloned gene sequences were confirmed via Sanger sequencing using vector and gene-specific primers and transformed into *E. coli* Rosetta BL21 (DE3) cells for heterologous expression.

Cloned ASAT1 in Rosetta BL21 (DE3) cells were used for protein expression and purification. 20mL culture was grown overnight at 37°C, shaking at 200 rpm in LB media supplemented with 1% filter sterilized glucose. Overnight culture was used to inoculate 3L of fresh LB media, which was incubated at 37°C shaking at 200rpm until an OD600 of 0.6 was reached. The cultures were then shocked on ice for 15 minutes, then induced with 0.4M IPTG. The cultures were then shaken at 18°C overnight. Cells were harvested by centrifugation at 5,400g for 10 minutes at 4°C. Pellets were resuspended in extraction buffer (25mM HEPES pH 7.5, 50mM KCl, 10% glycerol) and sonicated on ice for a total of 3 minutes, followed by a higher-speed centrifugation step for 30 minutes at 30,000g. The supernatant was used for purification using Ni-NTA Agarose (Qiagen). Protein was washed stepwise with 10mL each of increasing concentrations of imidazole (20mM, 40mM, and 60mM), then with a 5mL of a final wash at 500mM. The final wash was concentrated using 30,000 molecular weight cut-off centrifugal filters. Glycerol was added to a final concentration of about 50%, and then protein was stored at −20°C for subsequent analyses. Protein preps were analyzed by immunoblot analysis using a histidine-tag antibody (Supplementary Figure 17).

### ASAT in vitro characterization

For functional characterization, purified ASAT1s were subjected to *in vitro* enzymatic assays. Enzymatic assays consisted of 250µM commercially available acyl-CoAs of varying lengths, 2mM sucrose, 250mM sodium phosphate buffer pH 6, and 10µL recombinantly purified enzyme. Assays were incubated for one hour at 30°C and stopped by addition of 60µL 1:1 isopropanol and acetonitrile + 0.1% formic acid and 5µM telmisartan internal standard. Reactions were transferred to ice for 15 minutes, then centrifuged for 2 minutes. The supernatant was transferred to a glass vial with an insert and run on LC-MS equipped with a C18 column, with the same method used for acylsugar extracts (Supplementary Table 16). Activity was confirmed by the detection of the monoacylated sucrose using LC-MS. Specificity was quantified using peak area of the monoacylated product divided by the peak area of internal standard (telmisartan) and then normalized to the most abundant product. Two replicates of these assays were performed for each ASAT1.

## Acknowledgements

We thank the USDA Germplasm Resources Information Network (GRIN) for providing some of the *Physalis* seeds, the University of Missouri Genomics Core, and the University of Missouri Metabolomics core for assistance in sequencing and metabolite analyses. We are also indebted to Pilar Zamora-Tavares, Alan Herrera, Ofelia Vargas-Ponce, Mahinda Martínez, Luis Gerardo Hernandez-Sandoval, and Chelsea Pretz for their support during fieldwork in Mexico and for assistance with taxonomic identifications. We thank Elise Jensen and PiperJo Jones for their help with acylsugar sample preparation. We thank the staff of the herbaria CORD, H, IBUG, and QMEX for their help with curatorial tasks. This research was supported by the National Science Foundation (DEB 1902797 to SDS and DEB-2449574 to SDS and CAS). RD acknowledges the financial support for fieldwork and field extractions from the International Association for Plant Taxonomy (IAPT) through a Biodiversity Challenge grant, the Consejo Nactional de Investigaciones Científicas y Técnicas (CONICET, Argentina), the Secretaría de Ciencia y Tecnología de la Universidad Nacional de Córdoba (grant 203/14, SECYT-UNC, Argentina), and the European Union under the Maria Skłodowska-Curie grant agreement No 101151612 (MSCA).

## Data Availability

The code used in this study is available through GitHub https://github.com/lilliannowack. Physalis ASAT1s were deposited to NCBI with accession numbers *P. pruinosa* ASAT1 (PprASAT1): PV938311, *P. philadelphica* ASAT1 (PphASAT1): PV938309, and *P. coztomatl* ASAT1(PcoASAT1) PV938310.

## Author Contribution

LN conducted experiments, analyzed data, wrote and revised the manuscript. RD conducted experiments, analyzed data, and revised the manuscript. SDS planned the project, performed and analyzed data, and revised the manuscript. CAS planned and oversaw the project, analyzed data, and revised the manuscript.

## References

Barrientos L, Pérez-Castorena AL, Martínez M, and Maldonado E. Sucrose esters from the calyxes of *Physalis chenopodifolia*. Carbohydr Res. 2022:512:108518. 10.1016/j.carres.2022.108518

Beaudoin GAW and Facchini PJ. Benzylisoquinoline alkaloid biosynthesis in opium poppy. Planta. 2014:240(1):19–32. 10.1007/s00425-014-2056-8

Blomberg SP, Garland T, and Ives AR. Testing for phylogenetic signal in comparative data: behavioral traits are more labile. Evolution. 2003:57(4):717–745. 10.1111/j.0014-3820.2003.tb00285.x

Cao C-M, Wu X, Kindscher K, Xu L, and Timmermann BN. Withanolides and Sucrose Esters from *Physalis neomexicana*. J Nat Prod. 2015:78(10):2488–2493. 10.1021/acs.jnatprod.5b00698

Dale SM, Tomaszewski E, Lippman Z, and Van Eck J. Engineering the future of *Physalis grisea*: A focus on agricultural challenges, model species status, and applied improvements. PLANTS, PEOPLE, PLANET. 2024:6(6):1249–1260. 10.1002/ppp3.10536

D’Auria JC. Acyltransferases in plants: a good time to be BAHD. Curr Opin Plant Biol. 2006:9(3):331–340. 10.1016/j.pbi.2006.03.016

Deanna R, Larter MD, Barboza GE, and Smith SD. Repeated evolution of a morphological novelty: a phylogenetic analysis of the inflated fruiting calyx in the Physalideae tribe (Solanaceae). American Journal of Botany. 2019:106(2):270–279. 10.1002/ajb2.1242

Devi KP, Malar DS, Nabavi SF, Sureda A, Xiao J, Nabavi SM, and Daglia M. Kaempferol and inflammation: From chemistry to medicine. Pharmacological Research. 2015:99:1–10. 10.1016/j.phrs.2015.05.002

Erb M and Kliebenstein DJ. Plant secondary metabolites as defenses, regulators, and primary metabolites: The blurred functional trichotomy. Plant Physiol. 2020:184(1):39–52. 10.1104/pp.20.00433

Fan P, Miller AM, Liu X, Jones AD, and Last RL. Evolution of a flipped pathway creates metabolic innovation in tomato trichomes through BAHD enzyme promiscuity. Nat Commun. 2017:8(1):2080. 10.1038/s41467-017-02045-7

Fan P, Miller AM, Schilmiller AL, Liu X, Ofner I, Jones AD, Zamir D, and Last RL. In vitro reconstruction and analysis of evolutionary variation of the tomato acylsucrose metabolic network. Proc Natl Acad Sci USA. 2016:113(2):E239–248. 10.1073/pnas.1517930113

Fan P, Wang P, Lou Y-R, Leong BJ, Moore BM, Schenck CA, Combs R, Cao P, Brandizzi F, Shiu S-H, et al. Evolution of a plant gene cluster in Solanaceae and emergence of metabolic diversity. Elife. 2020:9:e56717. 10.7554/eLife.56717

Feng H, Acosta-Gamboa L, Kruse LH, Tracy JD, Chung SH, Nava Fereira AR, Shakir S, Xu H, Sunter G, Gore MA, et al. Acylsugars protect *Nicotiana benthamiana* against insect herbivory and desiccation. Plant Mol Biol. 2022:109(4):505–522. 10.1007/s11103-021-01191-3

Ferreira L, Vale A, de Souza A, Leite K, Sacramento C, Moreno M, Araújo T, Soares M, and Grassi M. Anatomical and phytochemical characterization of *Physalis angulata* L.: A Plant with Therapeutic Potential. Pharmacognosy Research. 2019:11(2):171–177. 10.4103/pr.pr_97_18

Fiesel PD, Kerwin RE, Jones AD, and Last RL. Trading acyls and swapping sugars: metabolic innovations in *Solanum* trichomes. Plant Physiol. 2024:196(2):1231–1253. 10.1093/plphys/kiae279

Fiesel PD, Parks HM, Last RL, and Barry CS. Fruity, sticky, stinky, spicy, bitter, addictive, and deadly: evolutionary signatures of metabolic complexity in the Solanaceae. Nat Prod Rep. 2022. 10.1039/D2NP00003B

Fritz SA and Purvis A. Selectivity in mammalian extinction risk and threat types: a new measure of phylogenetic signal strength in binary traits. Conserv Biol. 2010:24(4):1042– 1051. 10.1111/j.1523-1739.2010.01455.x

Gibson M, Santos WT, Oyler AR, Busta L, and Schenck CA. A new spin on chemotaxonomy using non-proteogenic amino acids as a test case. Applications in Plant Sciences. 2025. 10.1002/aps3.70006

Gu Z. Complex heatmap visualization. Imeta. 2022:1(3):e43. 10.1002/imt2.43

Gu Z, Eils R, and Schlesner M. Complex heatmaps reveal patterns and correlations in multidimensional genomic data. Bioinformatics. 2016:32(18):2847–2849. 10.1093/bioinformatics/btw313

Gu Z, Gu L, Eils R, Schlesner M, and Brors B. circlize Implements and enhances circular visualization in R. Bioinformatics. 2014:30(19):2811–2812. 10.1093/bioinformatics/btu393

He J, Alonge M, Ramakrishnan S, Benoit M, Soyk S, Reem NT, Hendelman A, Van Eck J, Schatz MC, and Lippman ZB. Establishing *Physalis* as a Solanaceae model system enables genetic reevaluation of the inflated calyx syndrome. Plant Cell. 2023:35(1):351–368. 10.1093/plcell/koac305

Huynh Nguyen K-N, Thi Nguyen N-V, and Ho Kim K. Determination of phenolic acids and flavonoids in leaves, calyces, and fruits of *Physalis angulata* L. in Viet Nam. Pharmacia. 2021:68(2):501–509. 10.3897/pharmacia.68.e66044

Jan R, Khan M, Asaf S, Lubna, Asif S, and Kim K-M. Bioactivity and Therapeutic Potential of Kaempferol and Quercetin: New Insights for Plant and Human Health. Plants. 2022:11(19):2623. 10.3390/plants11192623

Kasali FM, Tuyiringire N, Peter EL, Ahovegbe LY, Ali MS, Tusiimire J, Ogwang PE, Kadima JN, and Agaba AG. Chemical constituents and evidence-based pharmacological properties of *Physalis peruviana* L.: An overview. J Herbmed Pharmacol. 2021:11(1):35–47. 10.34172/jhp.2022.04

Kerwin RE, Hart JE, Fiesel PD, Lou Y-R, Fan P, Jones AD, and Last RL. Tomato root specialized metabolites evolved through gene duplication and regulatory divergence within a biosynthetic gene cluster. Sci Adv. 2024:10(17):eadn3991. 10.1126/sciadv.adn3991

Leckie BM, D’Ambrosio DA, Chappell TM, Halitschke R, De Jong DM, Kessler A, Kennedy GG, and Mutschler MA. Differential and synergistic functionality of acylsugars in suppressing oviposition by insect herbivores. PLoS One. 2016:11(4):e0153345. 10.1371/journal.pone.0153345

Li J, Song C, and He C. Chinese lantern in *Physalis* is an advantageous morphological novelty and improves plant fitness. Sci Rep. 2019:9(1):596. 10.1038/s41598-018-36436-7

Lopez-Gomollon S. *Physalis*: A new model crop to understand plant diversity. Plant Cell. 2023:35(1):338–339. 10.1093/plcell/koac315

Luu VT, Weinhold A, Ullah C, Dressel S, Schoettner M, Gase K, Gaquerel E, Xu S, and Baldwin IT. O-Acyl Sugars Protect a Wild Tobacco from Both Native Fungal Pathogens and a Specialist Herbivore. Plant Physiol. 2017:174(1):370–386. 10.1104/pp.16.01904

Lybrand DB, Anthony TM, Jones AD, and Last RL. An integrated analytical approach reveals trichome acylsugar metabolite diversity in the wild tomato *Solanum pennellii*. Metabolites. 2020:10(10). 10.3390/metabo10100401

Maldonado E, Torres FR, Martínez M, and Pérez-Castorena AL. Sucrose esters from the fruits of *Physalis nicandroides* var. attenuata. J Nat Prod. 2006:69(10):1511–1513. 10.1021/np060274l

Marone D, Mastrangelo AM, Borrelli GM, Mores A, Laidò G, Russo MA, and Ficco DBM. Specialized metabolites: Physiological and biochemical role in stress resistance, strategies to improve their accumulation, and new applications in crop breeding and management. Plant Physiol Biochem. 2022:172:48–55. 10.1016/j.plaphy.2021.12.037

Martínez M, Vargas-Ponce O, and Zamora-Tavares P. Taxonomic revision of *Physalis* in Mexico. Front Genet. 2023:14:1080176. 10.3389/fgene.2023.1080176

Medina-Medrano JR, Almaraz-Abarca N, González-Elizondo MS, Uribe-Soto JN, González-Valdez LS, and Herrera-Arrieta Y. Phenolic constituents and antioxidant properties of five wild species of *Physalis* (Solanaceae). Bot Stud. 2015:56:24. 10.1186/s40529-015-0101-y

Mierziak J, Kostyn K, and Kulma A. Flavonoids as important molecules of plant interactions with the environment. Molecules. 2014:19(10):16240–16265. 10.3390/molecules191016240

Moghe GD, Leong BJ, Hurney SM, Daniel Jones A, and Last RL. Evolutionary routes to biochemical innovation revealed by integrative analysis of a plant-defense related specialized metabolic pathway. Elife. 2017:6:e28468. 10.7554/eLife.28468

Nadakuduti SS, Uebler JB, Liu X, Jones AD, and Barry CS. Characterization of trichome-expressed BAHD acyltransferases in *Petunia axillaris* reveals distinct acylsugar assembly mechanisms within the Solanaceae. Plant Physiol. 2017:175(1):36–50. 10.1104/pp.17.00538

Orme D. The caper package: comparative analysis of phylogenetics and evolution in R. 2023.

Ovenden SPB, Yu J, Bernays J, Wan SS, Christophidis LJ, Sberna G, Tait RM, Wildman HG, Lebeller D, Lowther J, et al. Physaloside A, an acylated sucrose ester from *Physalis viscosa*. J Nat Prod. 2005:68(2):282–284. 10.1021/np049746r

Paradis E and Schliep K. ape 5.0: an environment for modern phylogenetics and evolutionary analyses in R. Bioinformatics. 2019:35(3):526–528. 10.1093/bioinformatics/bty633

Periferakis A, Periferakis K, Badarau IA, Petran EM, Popa DC, Caruntu A, Costache RS, Scheau C, Caruntu C, and Costache DO. Kaempferol: antimicrobial properties, sources, clinical, and traditional applications. International Journal of Molecular Sciences. 2022:23(23):15054. 10.3390/ijms232315054

Pichersky E and Lewinsohn E. Convergent evolution in plant specialized metabolism. Annu Rev Plant Biol. 2011:62:549–566. 10.1146/annurev-arplant-042110-103814

Popova V, Ivanova T, Stoyanova M, Mazova N, Dimitrova-Dyulgerova I, Stoyanova A, Ercisli S, Assouguem A, Kara M, Topcu H, et al. Phytochemical analysis of leaves and stems of *Physalis alkekengi* L. (Solanaceae). Open Chemistry. 2022:20(1):1292–1303. 10.1515/chem-2022-0226

Puterka GJ, Farone W, Palmer T, and Barrington A. Structure-function relationships affecting the insecticidal and miticidal activity of sugar esters. J Econ Entomol. 2003:96(3):636–644. 10.1603/0022-0493-96.3.636

Revell LJ. phytools 2.0: an updated R ecosystem for phylogenetic comparative methods (and other things). PeerJ. 2024:12:e16505. 10.7717/peerj.16505

Särkinen T, Bohs L, Olmstead RG, and Knapp S. A phylogenetic framework for evolutionary study of the nightshades (Solanaceae): a dated 1000-tip tree. BMC Evol Biol. 2013:13(1):214. 10.1186/1471-2148-13-214

Schenck CA, Anthony TM, Jacobs M, Jones AD, and Last RL. Natural variation meets synthetic biology: Promiscuous trichome-expressed acyltransferases from *Nicotiana*. Plant Physiol. 2022:190(1):146–164. 10.1093/plphys/kiac192

Schenck CA and Last RL. Location, location! cellular relocalization primes specialized metabolic diversification. FEBS J. 2020:287(7):1359–1368. 10.1111/febs.15097

Schenck CA, Westphal J, Jayaraman D, Garcia K, Wen J, Mysore KS, Ané J-M, Sumner LW, and Maeda HA. Role of cytosolic, tyrosine-insensitive prephenate dehydrogenase in *Medicago truncatula*. Plant Direct. 2020:4(5):e00218. 10.1002/pld3.218

Schilmiller AL, Charbonneau AL, and Last RL. Identification of a BAHD acetyltransferase that produces protective acyl sugars in tomato trichomes. Proc Natl Acad Sci U S A. 2012:109(40):16377–16382. 10.1073/pnas.1207906109

Shenstone E, Lippman Z, and Van Eck J. A review of nutritional properties and health benefits of Physalis species. Plant Foods Hum Nutr. 2020:75(3):316–325. 10.1007/s11130-020-00821-3

Shomali A, Das S, Arif N, Sarraf M, Zahra N, Yadav V, Aliniaeifard S, Kumar Chauhan D, and Hasanuzzaman M. Diverse physiological roles of flavonoids in plant environmental stress responses and tolerance. https://pmc.ncbi.nlm.nih.gov/articles/PMC9699315/. Retrieved July 15, 2025

Slocombe SP, Schauvinhold I, McQuinn RP, Besser K, Welsby NA, Harper A, Aziz N, Li Y, Larson TR, Giovannoni J, et al. Transcriptomic and reverse genetic analyses of branched-chain fatty acid and acyl sugar production in *Solanum pennellii* and *Nicotiana benthamiana*. Plant Physiol. 2008:148(4):1830–1846. 10.1104/pp.108.129510

de Souza AX, Riederer M, and Leide J. Multifunctional contribution of the inflated fruiting calyx: implication for cuticular barrier profiles of the Solanaceous genera *Physalis*, *Alkekengi*, and *Nicandra*. Front Plant Sci. 2022:13:888930. 10.3389/fpls.2022.888930

Tian C, Liu X, Chang Y, Wang R, Lv T, Cui C, and Liu M. Investigation of the anti-inflammatory and antioxidant activities of luteolin, kaempferol, apigenin and quercetin. South African Journal of Botany. 2021:137:257–264. 10.1016/j.sajb.2020.10.022

Tsugawa H, Cajka T, Kind T, Ma Y, Higgins B, Ikeda K, Kanazawa M, VanderGheynst J, Fiehn O, and Arita M. MS-DIAL: data-independent MS/MS deconvolution for comprehensive metabolome analysis. Nat Methods. 2015:12(6):523–526. 10.1038/nmeth.3393

Vellosa JCR, Regasini LO, Khalil NM, Bolzani V da S, Khalil OAK, Manente FA, Pasquini Netto H, and Oliveira OMM de F. Antioxidant and cytotoxic studies for kaempferol, quercetin and isoquercitrin. Eclet Quím. 2011:36:07–20. 10.1590/S0100-46702011000200001

Vendemiatti E, Hernández-De Lira IO, Snijders R, Torne-Srivastava T, Therezan R, Simioni Prants G, Lopez-Ortiz C, Reddy UK, Bleeker P, Schenck CA, et al. *Woolly* mutation with the *Get02* locus overcomes the polygenic nature of trichome-based pest resistance in tomato. Plant Physiol. 2024a:195(2):911–923. 10.1093/plphys/kiae128

Vendemiatti E, Nowack L, Peres LEP, Benedito VA, and Schenck CA. Sticky business: the intricacies of acylsugar biosynthesis in the Solanaceae. Phytochem Rev. 2024b. 10.1007/s11101-024-09996-y

Voo SS, Grimes HD, and Lange BM. Assessing the biosynthetic capabilities of secretory glands in Citrus peel. Plant Physiol. 2012:159(1):81–94. 10.1104/pp.112.194233

Wang H, Ren J, Zhou S, Duan Y, Zhu C, Chen C, Liu Z, Zheng Q, Xiang S, Xie Z, et al. Molecular regulation of oil gland development and biosynthesis of essential oils in *Citrus spp*. Science. 2024:383(6683):659–666. 10.1126/science.adl2953

Weinhold A and Baldwin IT. Trichome-derived O-acyl sugars are a first meal for caterpillars that tags them for predation. Proc Natl Acad Sci U S A. 2011:108(19):7855–7859. 10.1073/pnas.1101306108

Wen X, Hempel J, Schweiggert RM, Ni Y, and Carle R. Carotenoids and Carotenoid Esters of Red and Yellow *Physalis* (*Physalis alkekengi* L. and *P. pubescens* L.) Fruits and Calyces. J Agric Food Chem. 2017:65(30):6140–6151. 10.1021/acs.jafc.7b02514

Weng J-K. The evolutionary paths towards complexity: a metabolic perspective. New Phytol. 2014:201(4):1141–1149. 10.1111/nph.12416

Weng J-K, Philippe RN, and Noel JP. The rise of chemodiversity in plants. Science. 2012:336(6089):1667–1670. 10.1126/science.1217411

Wickham H. ggplot2: Elegant Graphics for Data Analysis (Springer: New York, NY). 10.1007/978-0-387-98141-3

Wilf P, Carvalho MR, Gandolfo MA, and Cúneo NR. Eocene lantern fruits from Gondwanan Patagonia and the early origins of Solanaceae. Science. 2017:355(6320):71–75. 10.1126/science.aag2737

Wu M, Northen TR, and Ding Y. Stressing the importance of plant specialized metabolites: omics-based approaches for discovering specialized metabolism in plant stress responses. Front Plant Sci. 2023:14:1272363. 10.3389/fpls.2023.1272363

Xu D, Wang Z, Zhuang W, Wang T, and Xie Y. Family characteristics, phylogenetic reconstruction, and potential applications of the plant BAHD acyltransferase family. Front Plant Sci. 2023:14:1218914. 10.3389/fpls.2023.1218914

Yuan X, Smith NSS, and Moghe GD. Analysis of plant metabolomics data using identification-free approaches. Applications in Plant Sciences. 2025:e70001. 10.1002/aps3.70001

Zhang C-R, Khan W, Bakht J, and Nair MG. New antiinflammatory sucrose esters in the natural sticky coating of tomatillo (*Physalis philadelphica*), an important culinary fruit. Food Chem. 2016:196:726–732. 10.1016/j.foodchem.2015.10.007

