## Supplementary Figures for "Novel production of structurally diverse and sticky defense metabolites on wild tomatillo fruits"

[6]Corresponding author

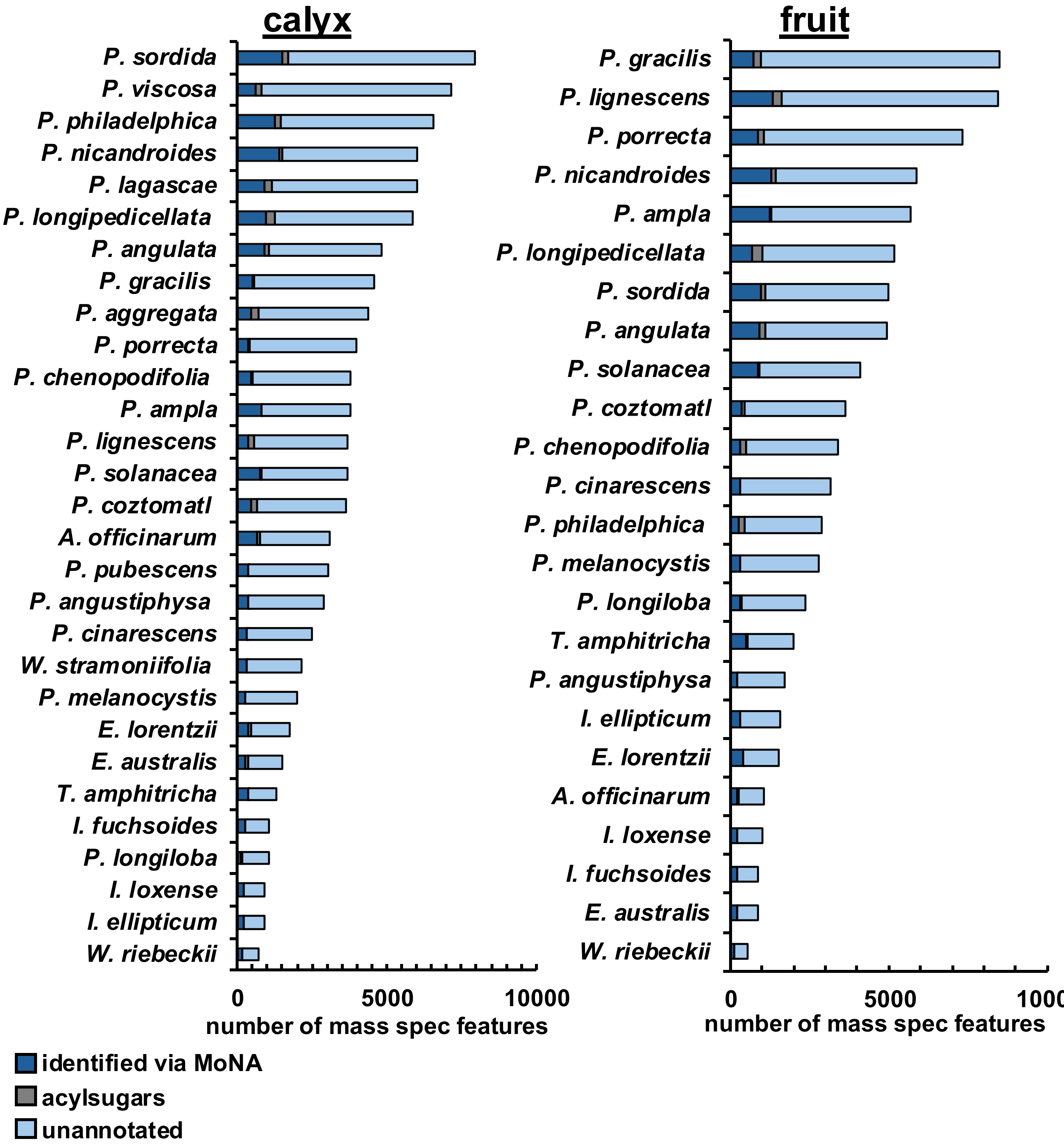


**Supplementary Figure 1 – Total numbers of mass spec features identified from fruit and calyx extracts.** Mass spec features from calyx and fruit extracts were identified using MSDial. Mass features were annotated using the Massbank of North America (MoNA) database (dark blue) with many features remaining unannotated (light blue). Acylsugars make up a small portion of all mass spec features identified (grey). Acylsugars are a combination of manually annotated acylsugars and putative acylsugars based on *m/z* of formate adducts (639 *m/z* to 835 *m/z*), retention time (6.0 minutes to 15.5 minutes), and presence of a formate adduct. Unannotated mass spec features make up most of the chemical diversity identified due to limitations in metabolite libraries and challenges with identification of mass spec features.

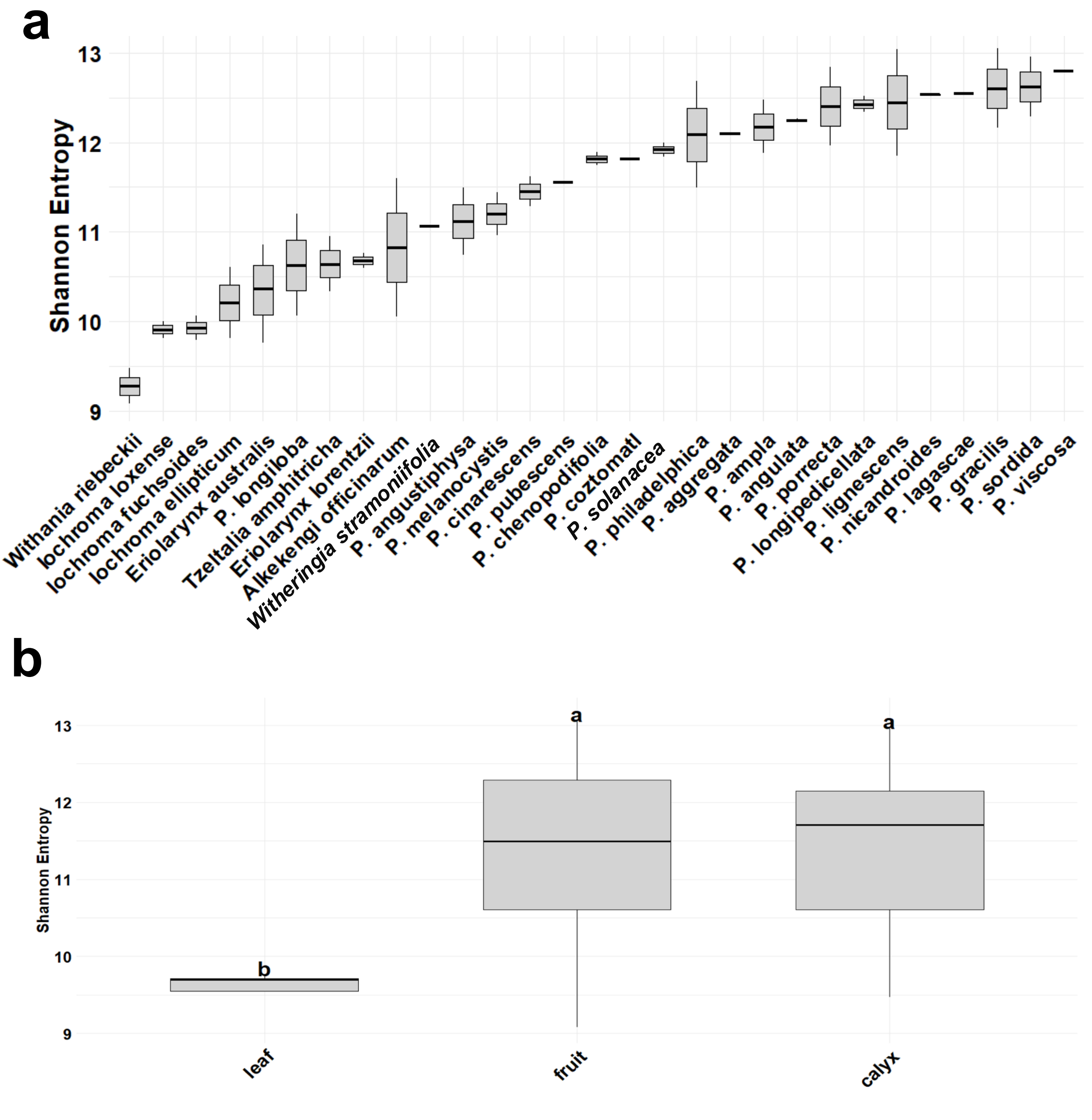


**Supplementary Figure 2 – Shannon entropy values per each species based on all identified mass spec features. (a)** Shannon entropy values were calculated for each tissue extract from all mass spec features identified and averaged to obtain a species average. Species with a flat line are represented by only on tissue extract. Peak area and height for mass spec features were not considered. **(b)** Shannon entropy values were averaged to obtain a tissue average across all species; individual points are not plotted. Sample sizes ($n$) were: leaf = 5, fruit = 25, calyx = 30. One-way ANOVA was performed to compare Shannon entropy values of different tissues. This revealed that there was a significant difference among tissues ($F(2) = 7.49$, $p = 0.0013$). Tukey post-hoc test showed that there were significant differences between fruit and leaf ($p = 0.0017$), and calyx and leaf ($p = 0.0011$), but not between calyx and fruit ($p = 0.9861$). Tissues not sharing a letter are significantly different.

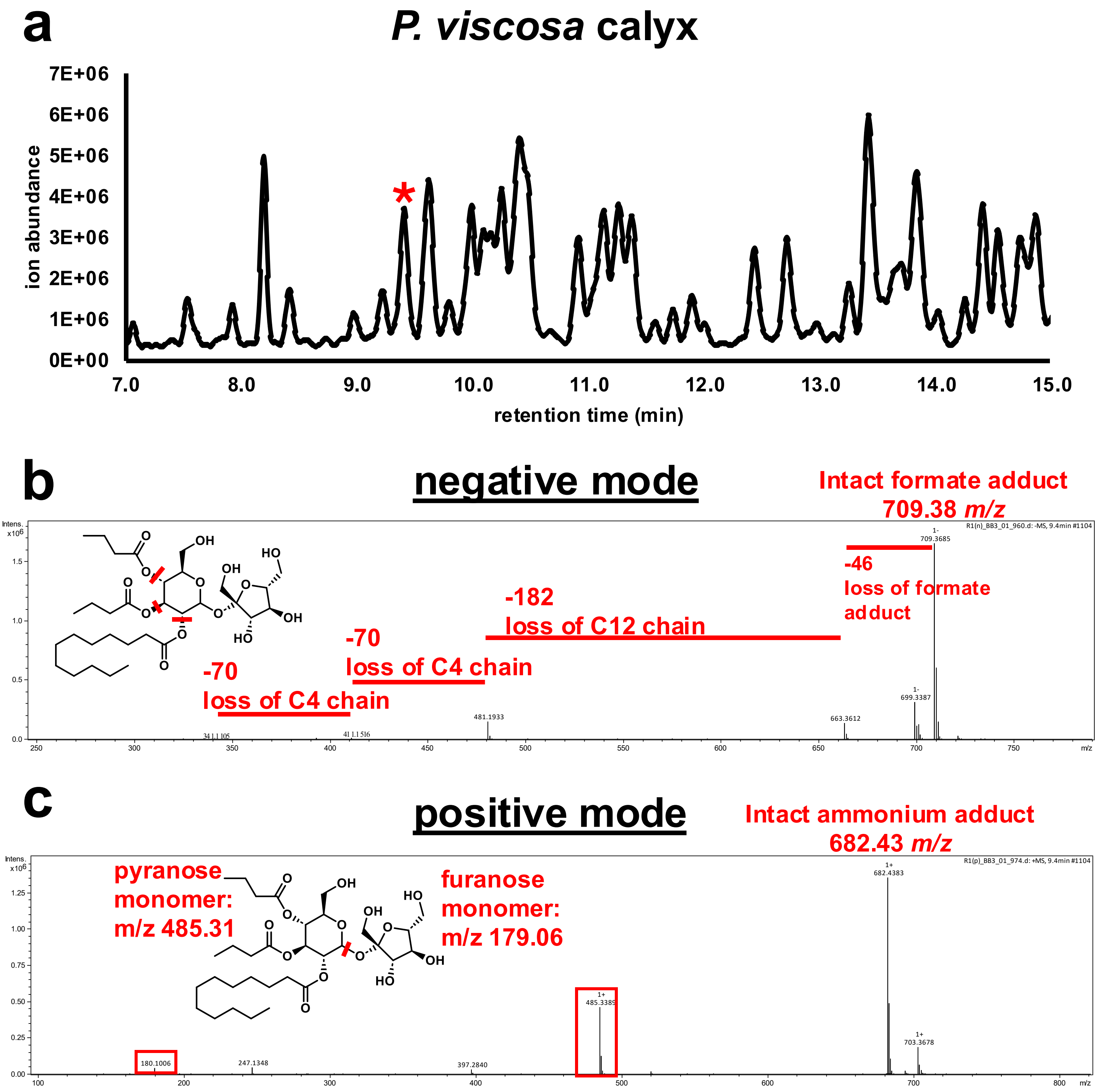


**Supplementary Figure 3 – Acylsugar annotations based on fragmentation pattern in negative and positive mode. (a)** Representative LC-MS chromatogram showing the range in which acylsugars elute. Red star indicates an acylsugar selected to demonstrate fragmentation patterns in **(c)** positive and **(b)** negative modes. **(b)** Negative mode acylsugar annotation with fragment ions and corresponding losses labeled in red. Positions of cleavage are shown in red. Neutral loss fragments indicate acyl chains of 4, 4, and 12 carbons in length. Overall acylsugar annotation S3(20):4,4,12, location of acyl chains on specific hydroxyl groups is unknown. **(c)** Positive mode acylsugar annotation with fragment ions for the pyranose and furanose portion. Fragmentation of the glycosidic linkage is shown in red on the structure yielding the two ions that are boxed in red. Fragment ions for this acylsugar show that all acyl chains are on the same ring (likely pyranose as drawn).

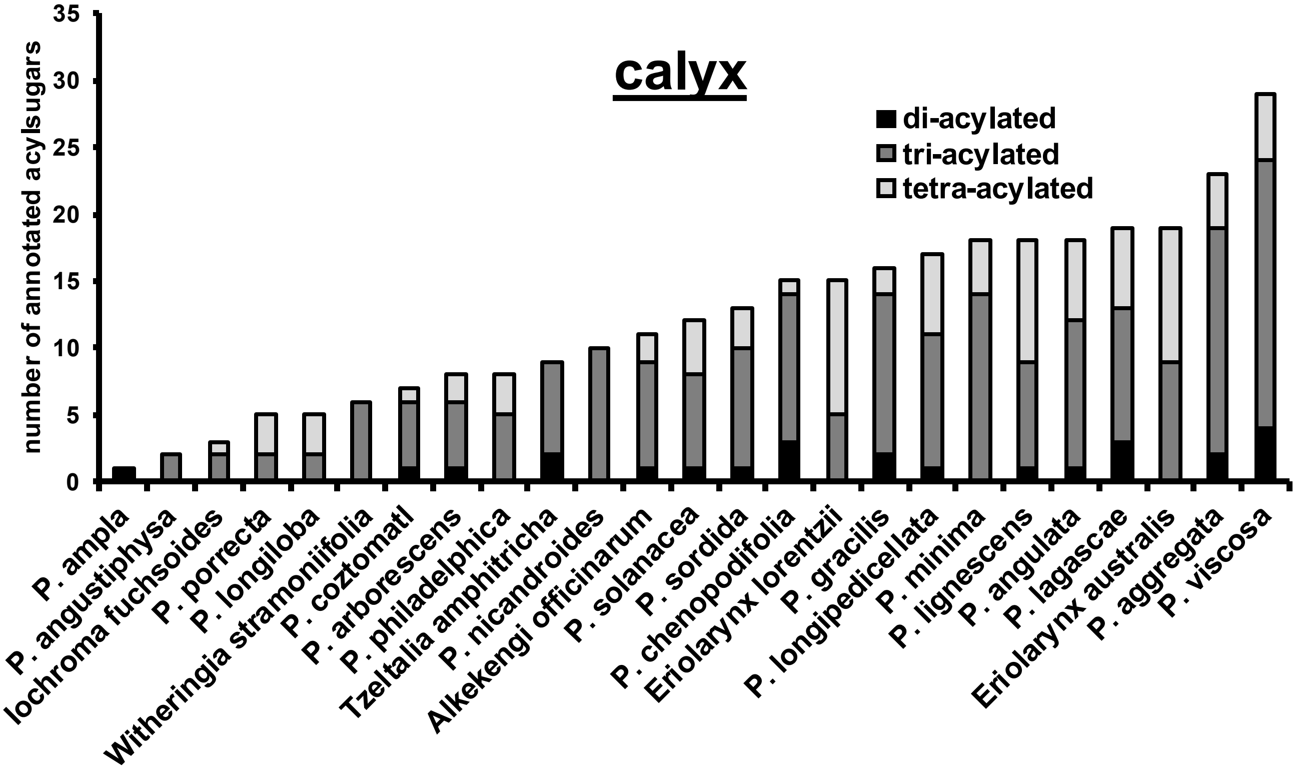


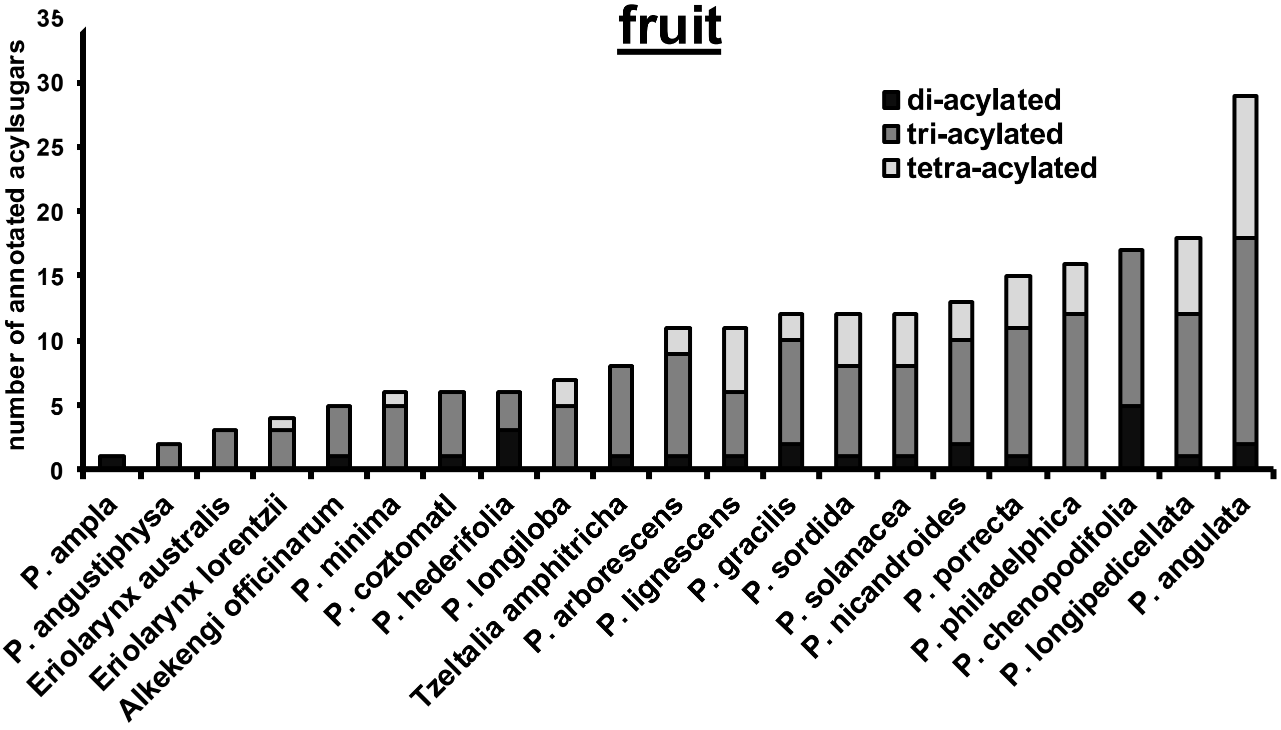


**Supplementary Figure 4 – Number of di, tri, and tetra acylated acylsugars in each species and tissue.** Annotated acylsugars were counted for each species and tissue and grouped by number of acylations. Triacylsugars are the most common across tissues and species with almost all species accumulating triacylsugars, whereas di and tetraacylsugars are less frequent. Not all species have both fruit and calyx extracts given the nature of the sampling; thus, the two graphs consist of slightly different numbers of species.

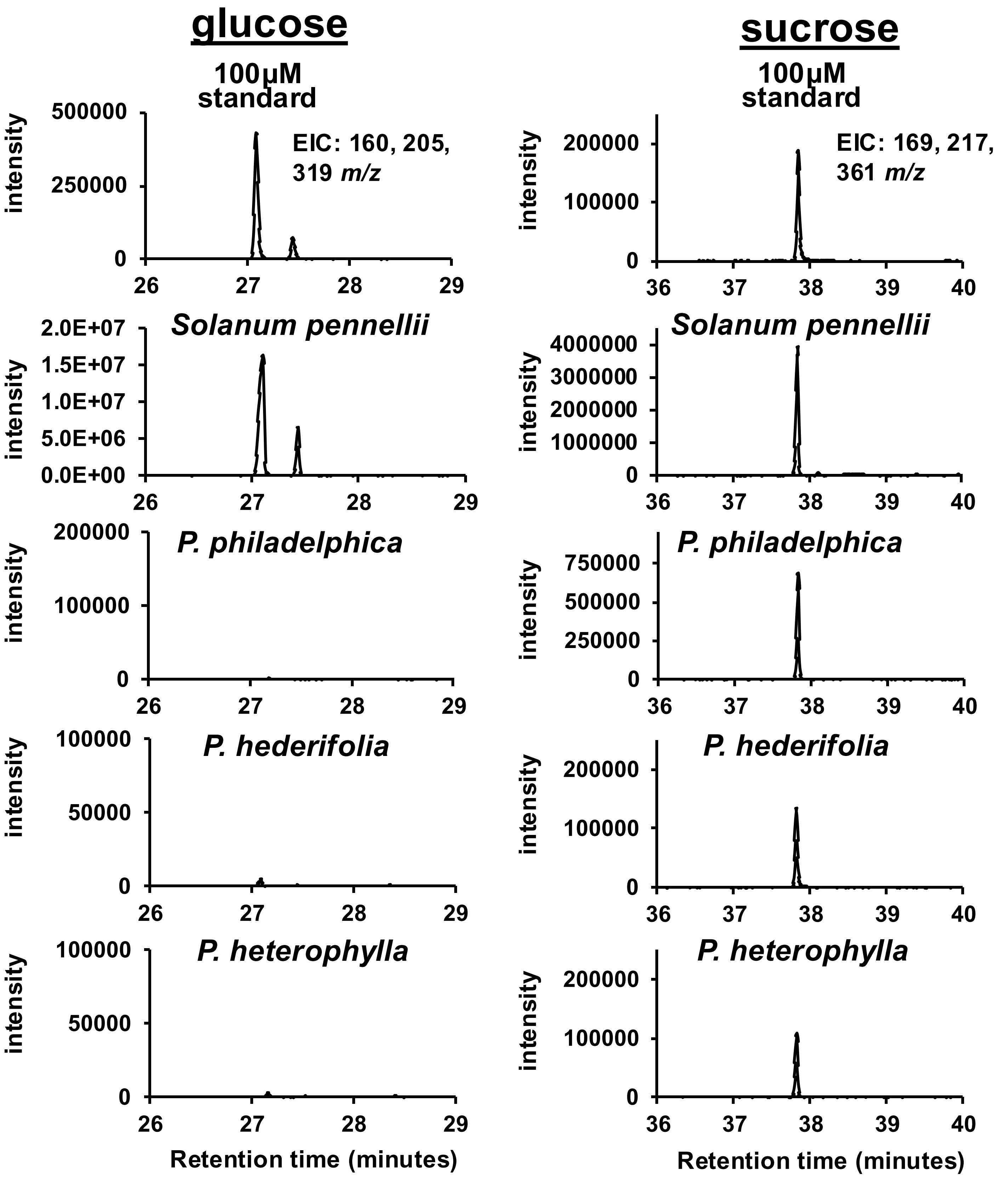


**Supplementary Figure 5 – GC-MS sugar core analysis of *Physalis* acylsugars.** Sugar cores were analyzed by extracting acylsugars, separating acylated sugars from free sugars, cleaving acyl chains from the acylated sugars, and derivatizing the remaining free sugars. Derivatized sugar cores and two derivatized sugar standards commonly found as the sugar cores of acylsugars (glucose (left) and sucrose (right)) were separated and identified using GC-MS. Extracted ion chromatograms (EIC) are shown for each standard and acylsugar extract at the given *m/z* values.

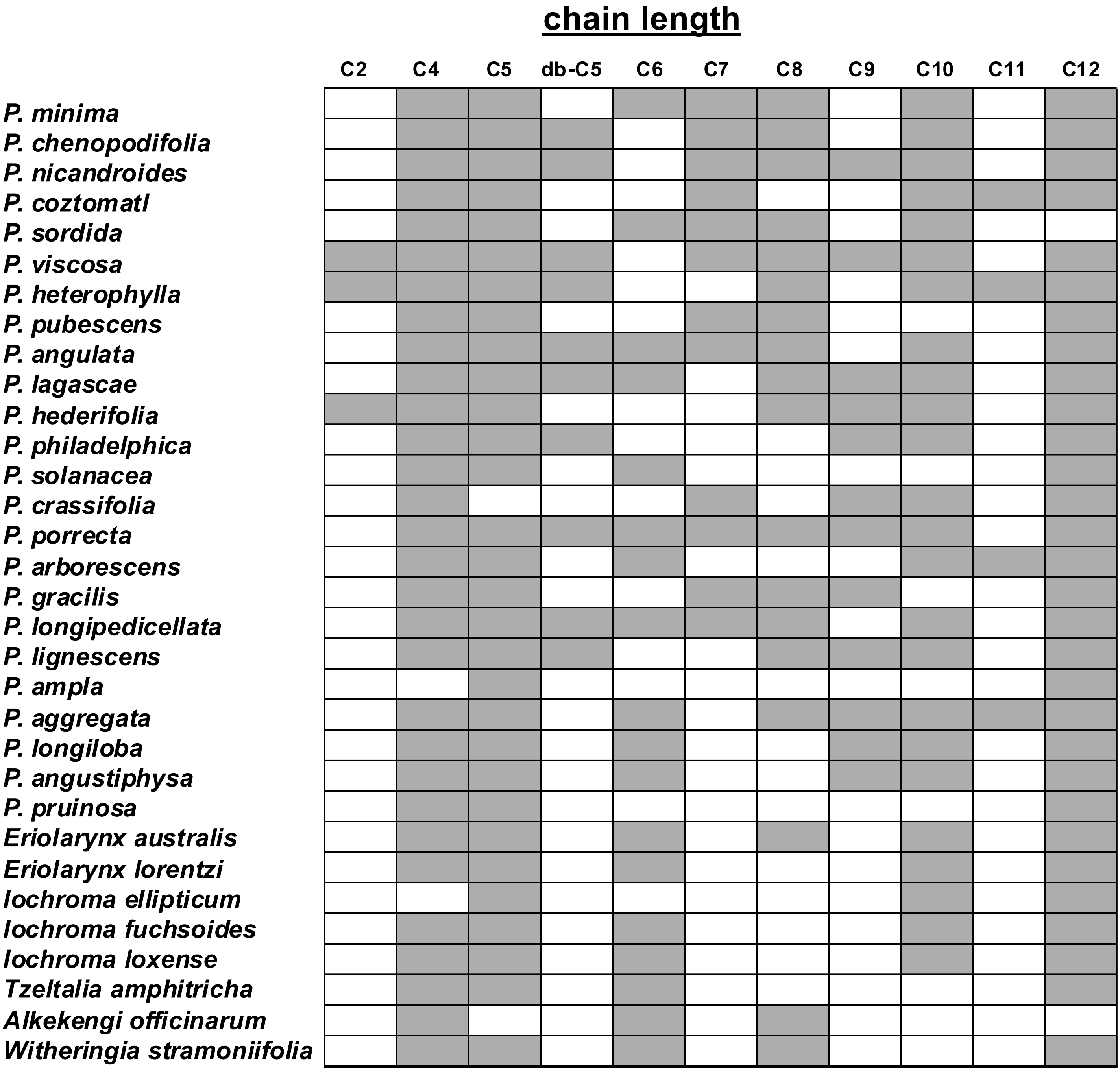


**Supplementary Figure 6 – Distribution of acyl chains determined from LC-MS-based acylsugar annotations.** Grey indicates presence and white indicates absence. Acyl chain length of C2 indicates an acetyl group with two carbons, C4 indicates an acyl chain with 4 carbons. db-C5 indicates an acyl chain with a likely double bond due to an observed 2 *m/z* difference from the C5 chain. Short chains (C4 and C5) as well as the medium chain C12 were the most observed acyl chains.

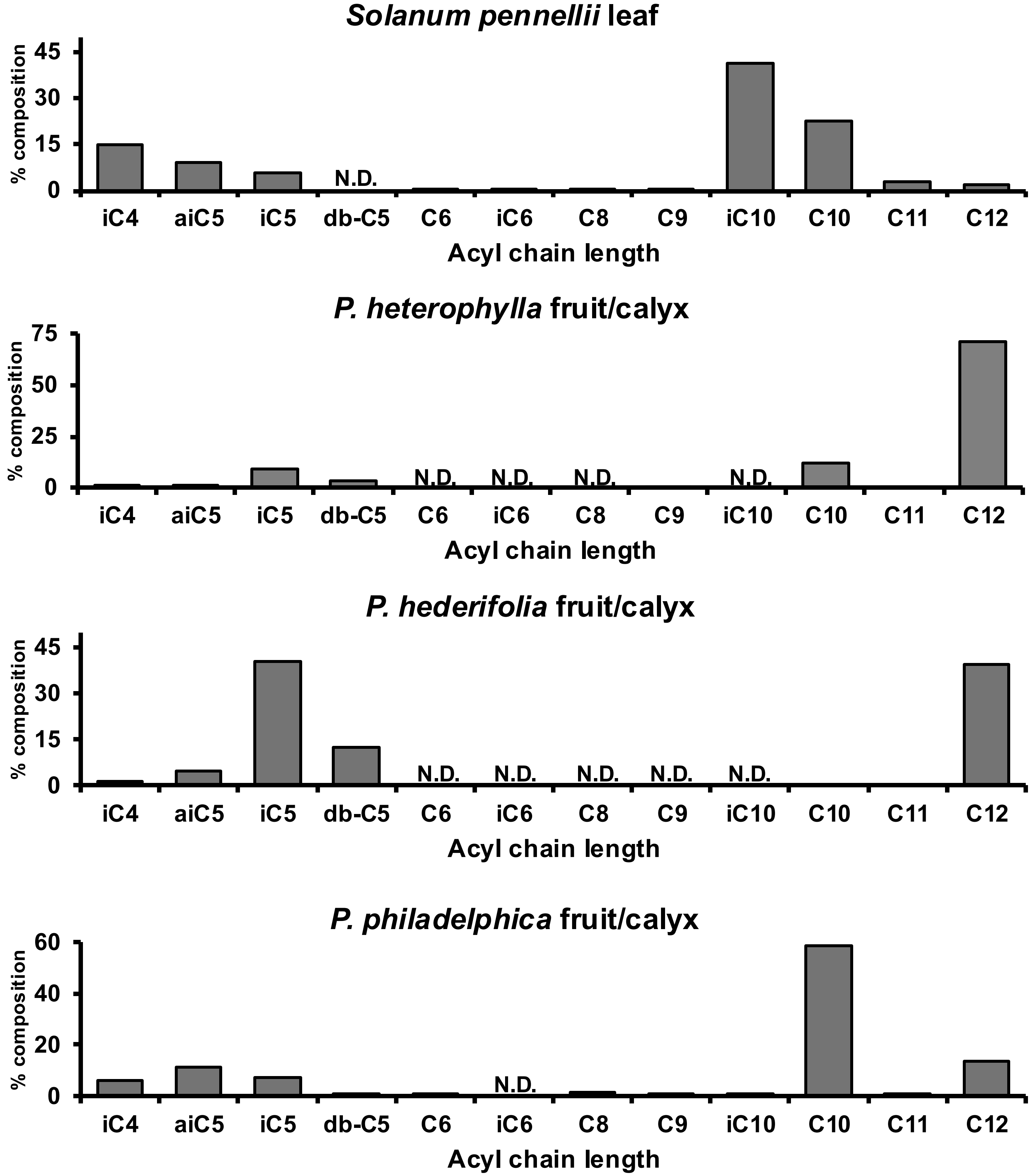


**Supplementary Figure 7 – GC-MS acyl chain analysis from a subset of *Physalis* species.** Acyl chains were prepared by cleaving acyl chains from acylsugar extracts and converting them to ethyl esters. Acyl chain ethyl esters were identified based on ethyl ester standards mix and NIST library searches. Percent composition was calculated using peak area for each chain length per peak area of all acyl chains. db indicates a double bond. Branched acyl chains include isobutyryl (iC4), isovaleryl (iC5), anteiso-valeryl (aiC5), isohexanoyl (iC6), and isodecanoic acid (iC10). GC-MS acyl chain analysis generally confirm LC-MS acylsugar annotations.

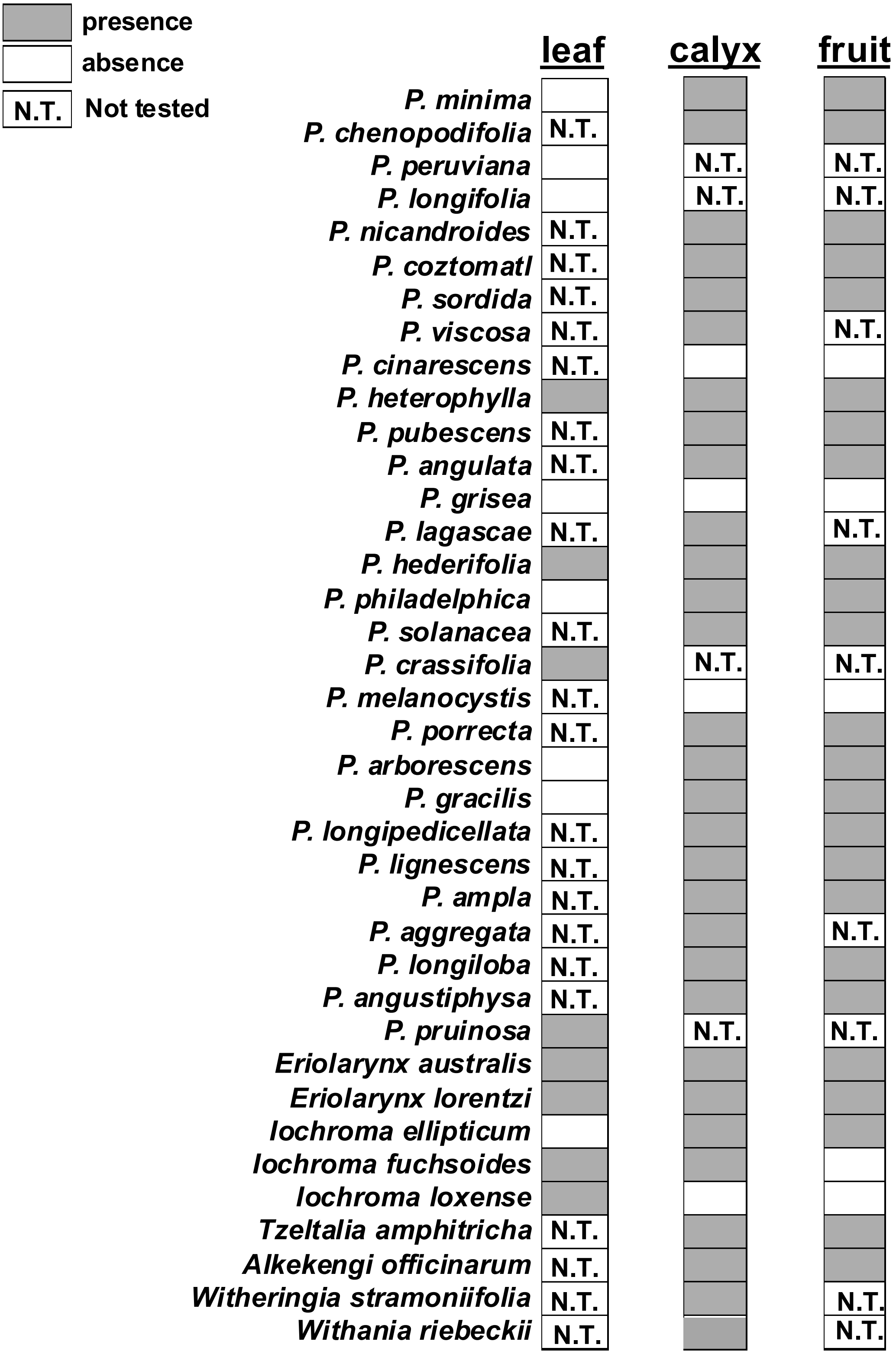


**Supplementary Figure 8 – Acylsugar absence and presence across all tissues and species.** Absence and presence was determined from LC-MS surface extracts. Absence indicates extracts for which acylsugars were not detected; however, it could be that they are below our detection limit and are present at very low abundances.

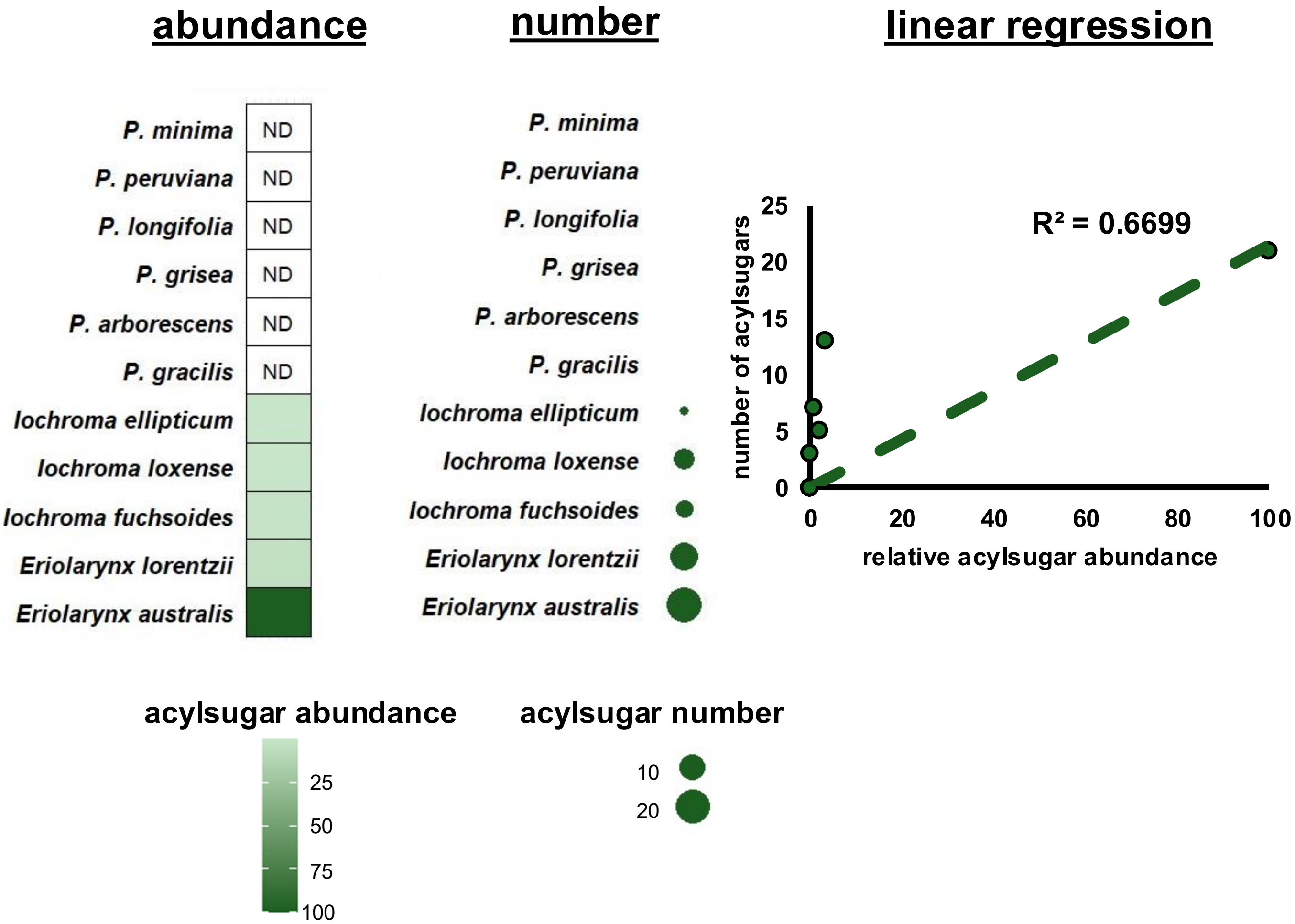


**Supplementary Figure 9 – Number and abundance of *Physalis* leaf acylsugars.** Acylsugars were quantified using the total peak area of all annotated acylsugars, corrected for internal standard and dry weight, and normalized to the most abundant extract of that particular tissues, in this case leaf. ND indicates species in which acylsugars were below detection limits of the LC-MS used. For number of acylsugars, the size of the circle indicates the number, with no circle indicating no detectable acylsugars. Linear regression of the relationship between relative abundance and number of acylsugars, with the $R^2$ value for the line of best fit shown.

a

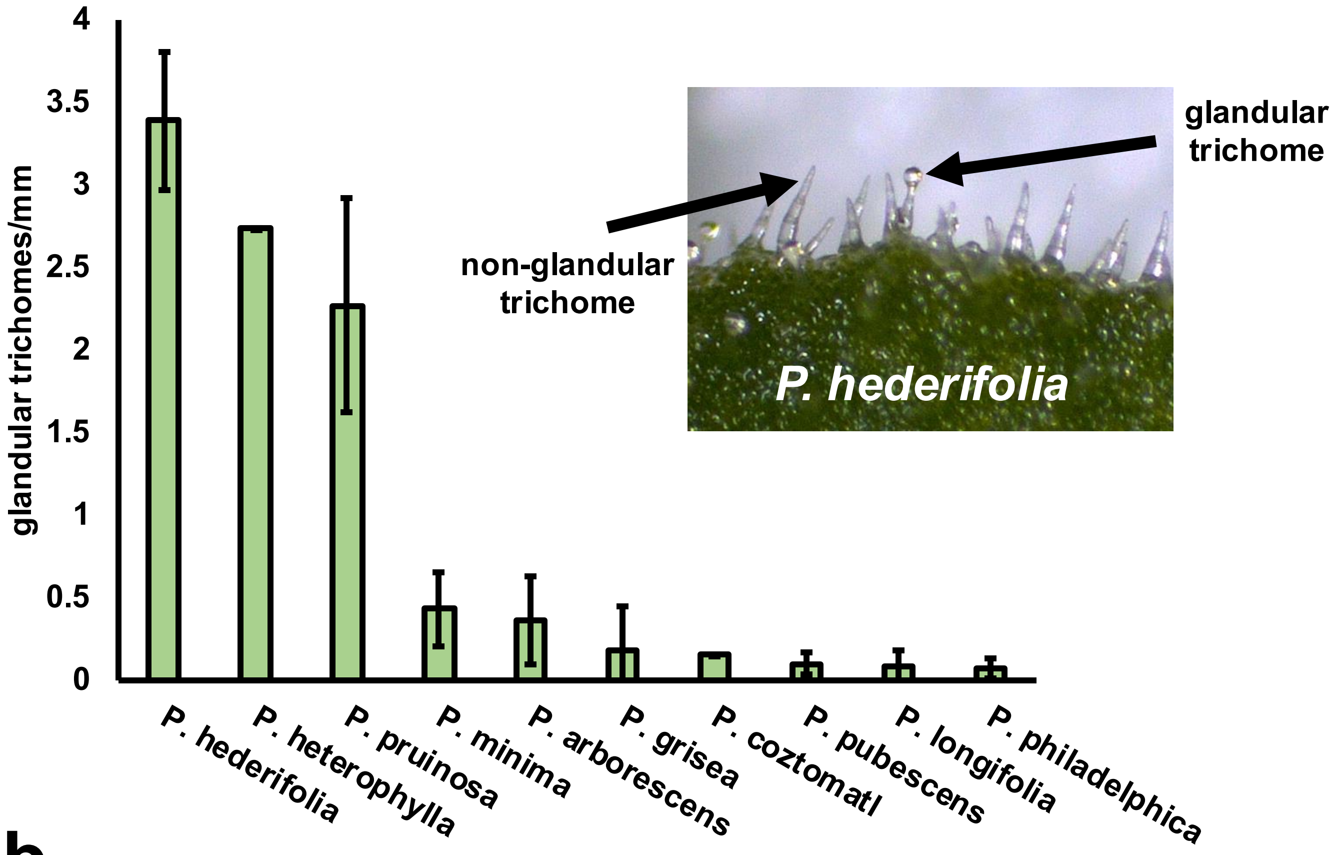


b

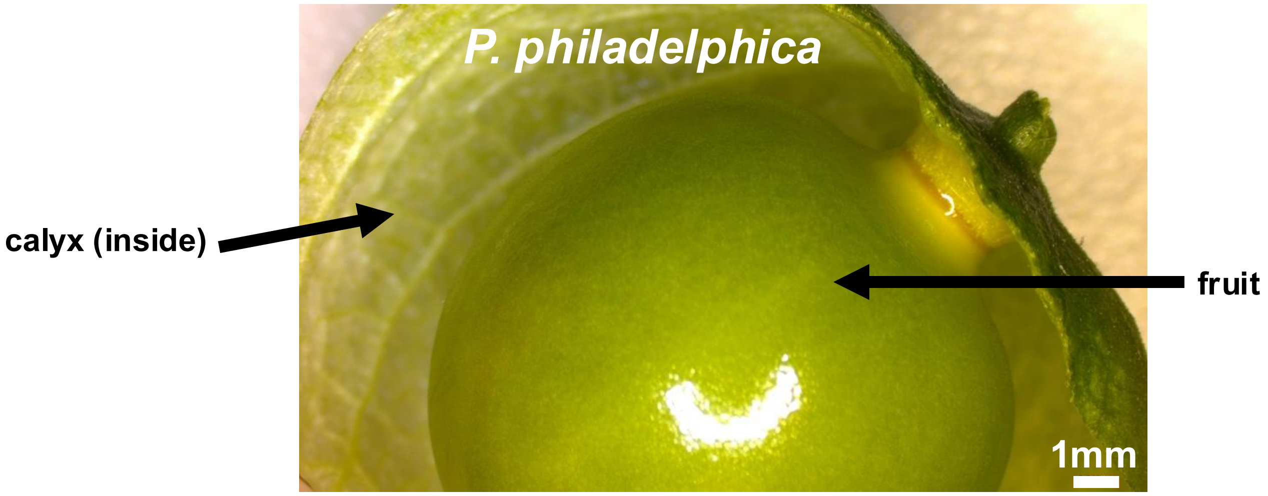


**Supplementary Figure 10 – *Physalis* glandular trichome abundance. (a)** Glandular trichomes were counted from three individual leaves. Glandular trichomes were identified as shown in the inset. Bars indicate number of trichomes per mm of leaf distance ± standard deviation. **(b)** Light microscopy image of *P. philadelphica* fruit surface and inside of calyx with no glandular trichomes.

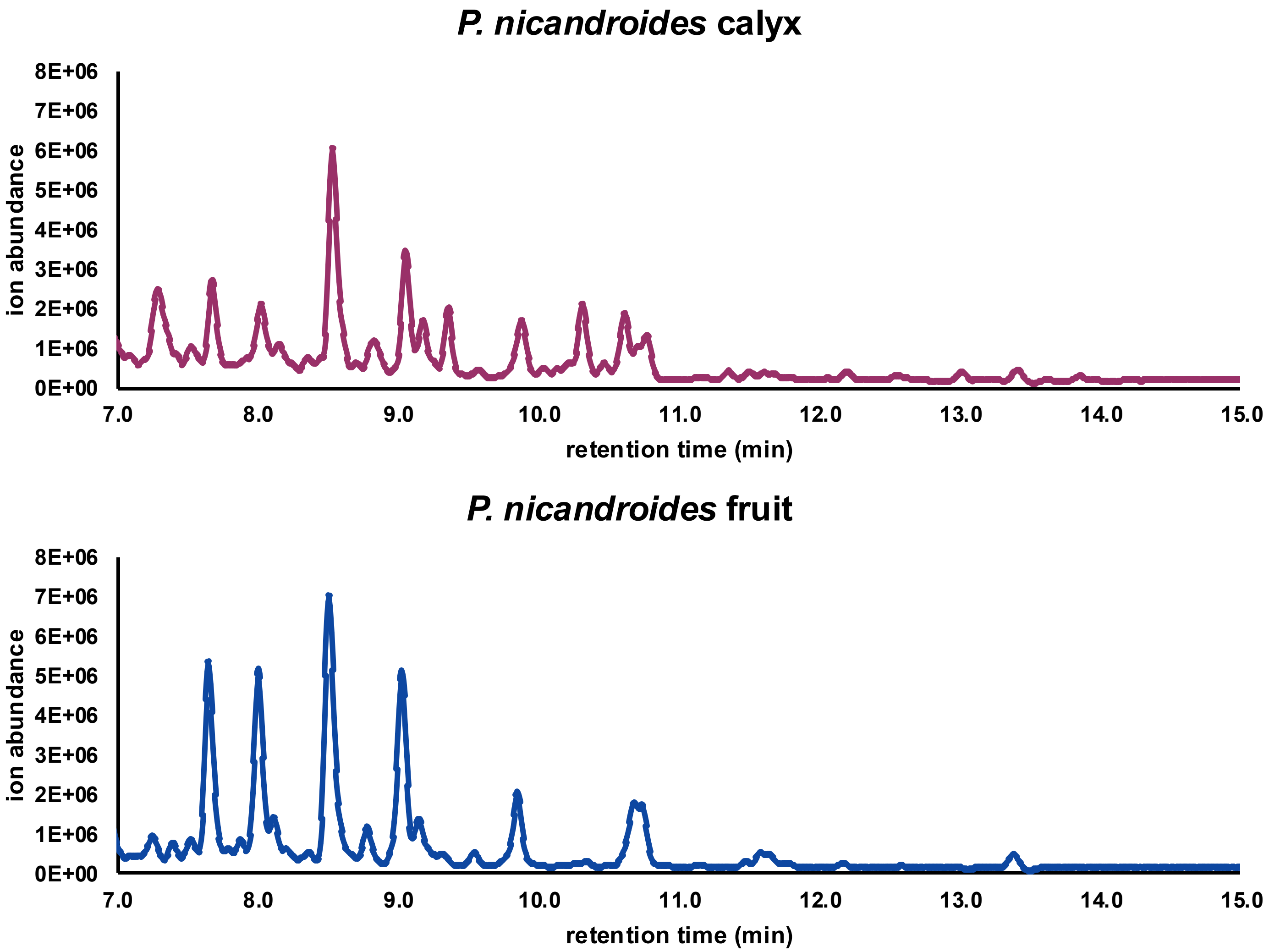


**Supplementary Figure 11 – *P. nicandroides* calyx and fruit extracts showing acylsugar retention time window.** Calyx and fruit surface extracts were separated using LC-MS. Total ion chromatogram for the retention time range that has abundant acylsugars is shown. In general, the same peaks are present in leaf and calyx extracts, however abundances vary and are greater in fruit extracts.

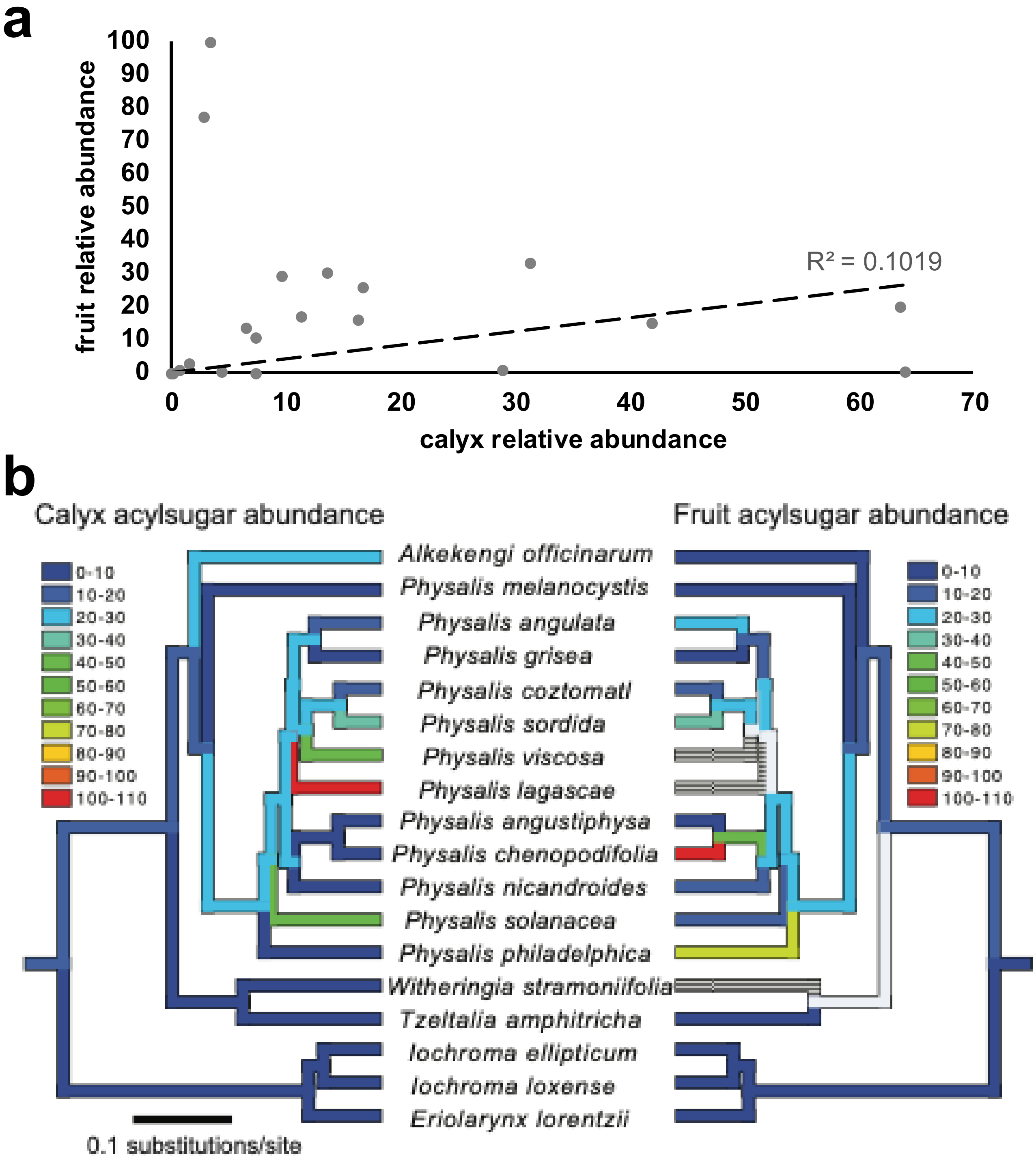


**Supplementary Figure 12 – Lack of correlation between fruit and calyx acylsugar abundance. (a)** Linear regression of relationship between relative abundance of fruit and calyx acylsugars. Only species in which both tissues were quantified were included. $R^2$ values of 0.1019 shows a lack of correlation between these two traits. **(b)** Fruit and calyx acylsugar abundances mapped across species included in the phylogeny of Deanna et al. (2019). The tree (from Deanna et al. 2019) was pruned to the taxa for which we have AS abundance data. Branches are colored by the abundance. Gray ancestral branches denote uncertainty, and gray striped branches correspond to missing tip data. Branch lengths are in units of substitutions per site. The relationship between these two traits were assessed with a phylogenetic generalized least squares analysis, including the alpha parameter to estimate the strength of phylogenetic signal. As in **(a)**, the two variables are not significantly related (P=0.79-0.99 across a sample of 100 phylogenetic trees from Deanna et al, 2019).

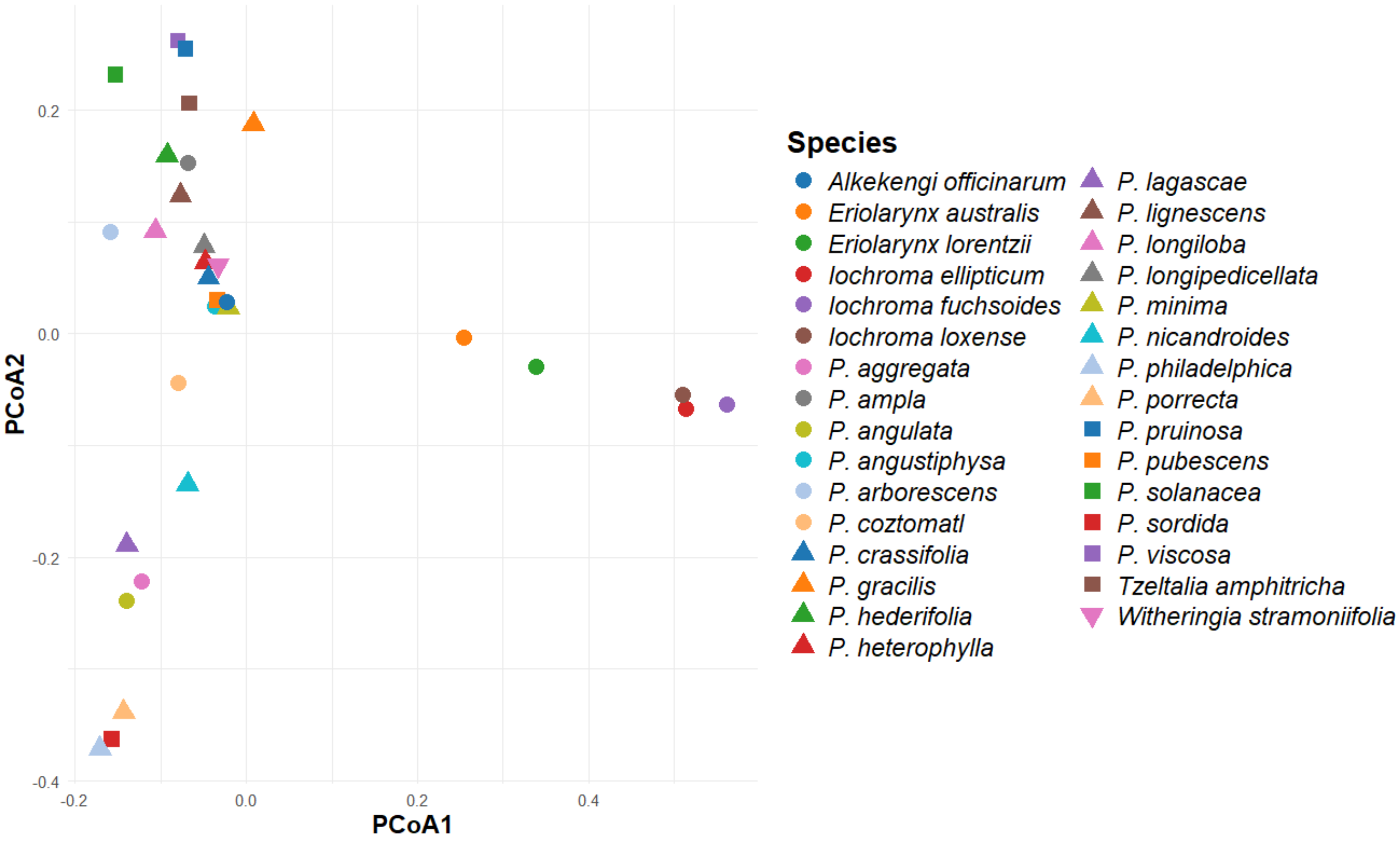


**Supplementary Figure 13 – Principal Coordinate Analysis (PCoA) of acylsugar diversity in *Physalis* and related species.** PCoA based on absence and presence of all annotated acylsugars. Acylsugars present in multiple tissues of the same species with identical *m/z,* fragmentation patterns, and retention time were considered one acylsugar.

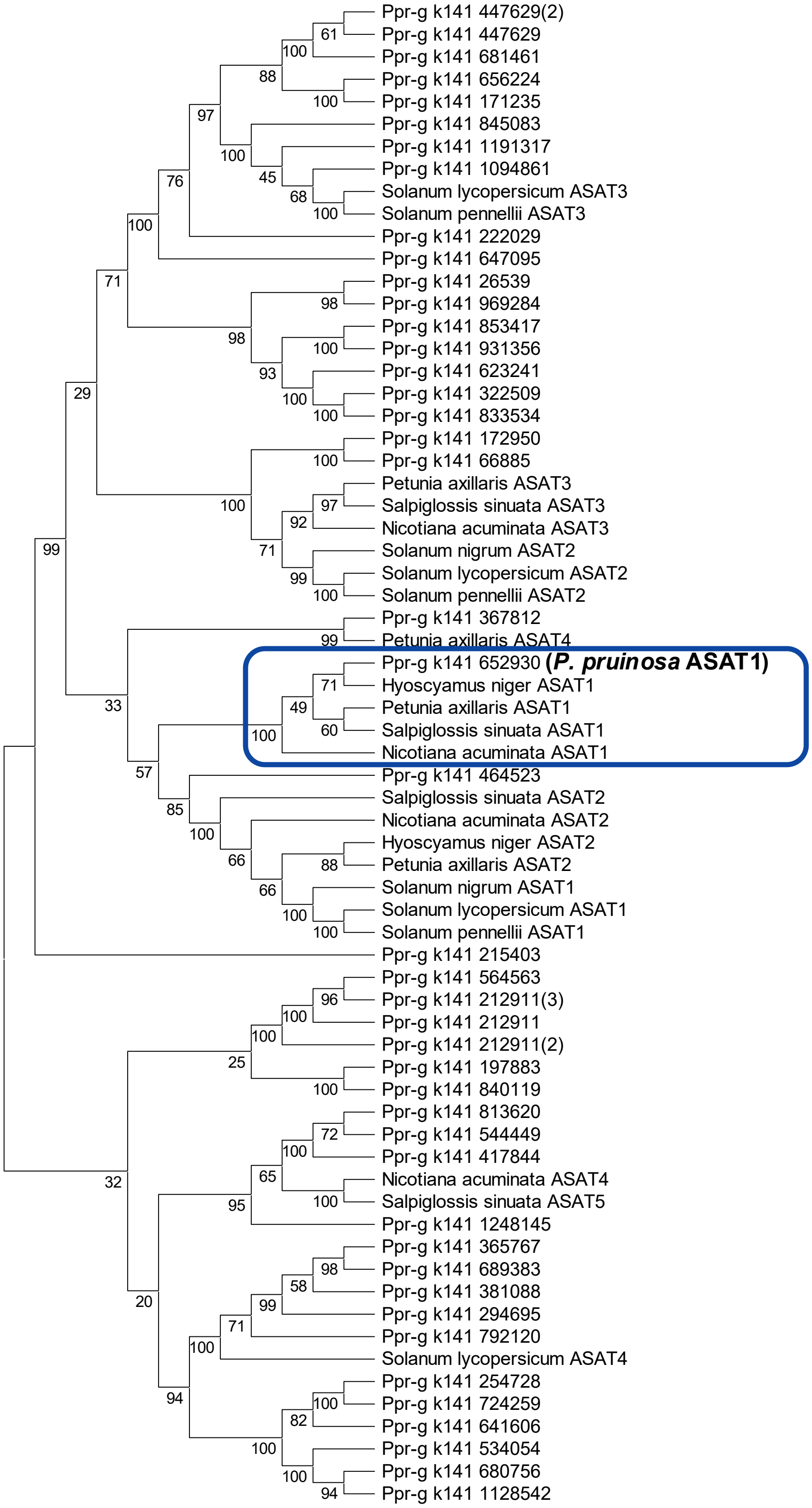


**Supplementary Figure 14 – Phylogeny of *P. pruinosa* ASATs together with biochemically characterized ASATs from other Solanaceae species.** ASAT candidates were identified in *P. pruinosa* from BLAST searches with tomato ASATs. Candidates were filtered based on amino acid length and presence of conserved substrate binding and catalytic motifs. Amino acid alignments were performed using MUSCLE and were used for phylogenetic analysis using the neighbor-joining method in MEGA12. 500 bootstrap replicates were calculated and are labeled above nodes. Ppr-g k141 indicates *P. pruinosa* sequences, all others are labeled accordingly. Blue box indicates ancestral ASAT1 clade, which acylate at the R2 position of sucrose.

```
Physalis_coztomatl_ASAT1         MAASALVSLSQKIIKPFSPTPLSERIYKLSFIDQFNSTQYVPVTLFYPNT
Physalis_philadelphica_ASAT1     MAASALVSLSKKIIKPFSPTPLSERIYKLSFIDQFNSTQYVPVTLFYPYT
Physalis_pruinosa_ASAT1          MAASALVSLSKKIIKPFSPTPFSERIYKLSFIDQFNSTQYVPVSLFYPNT
                                 **********:**********:*********************:**** *

Physalis_coztomatl_ASAT1         KGEPSVEPNDMCKVIENSLSEALASYYPFAGTLRDNVHIECNDIGADFYK
Physalis_philadelphica_ASAT1     KGEPSVEPNDMCKVIENSLSKALAAYYPFAGTLSDNVHIECNDIGADFYK
Physalis_pruinosa_ASAT1          KGEPSVEPNDMCKVIENSLSKALASYYPFAGTLRDNVHIECNDIGADFYM
                                 ********************:***:******** *************** 

Physalis_coztomatl_ASAT1         ARFDCPMSEILKSHDRNVKEMVYPKDIPWNVVGPDGKLVTVQLNQFDCGG
Physalis_philadelphica_ASAT1     ARFDCPMSEILKSHDRNVKEMVYPKDIPWNVVGPDGKLVTVQLNQFDCGG
Physalis_pruinosa_ASAT1          ARFDCPMSEILKSHDRNFKEMLYPKDIPWNVVGPNKKLVMVQFNQFDCGG
                                 *****************.***:************: *** **:*******

Physalis_coztomatl_ASAT1         IALSTCVTHKVADMCSMFKFIRDWASIARDSNSNIRPQFVGSSYFPPKNE
Physalis_philadelphica_ASAT1     IALSTCVTHKVADMCSMFKFIRDWASIARDSNSNIRPQFVGSSYFPPKNE
Physalis_pruinosa_ASAT1          IALSTCVTHKVADMCTMFKFVRDWASVARDSNFNIRPQFVGSSYFPPKNE
                                 ***************:****:*****:***** *****************

Physalis_coztomatl_ASAT1         PVNEPPREQCVTRRLAFSNRTLKSFISECSVPGVEKPSRIETLTALFYQC
Physalis_philadelphica_ASAT1     PVNEPPREQCVTRRLAFSNRTLKSFISECSVSGVEKPSRVETLTALFYKC
Physalis_pruinosa_ASAT1          PANEPPREQCVTKRLAFSNRTLKSFISECSVSGVEKPSRVETLTALFYKC
                                 *.**********:******************.*******:********:*

Physalis_coztomatl_ASAT1         GMRANSSDRLLKPSILFQTMNLRPFIPLPEDAAGNFSSSLSVPTYVEEEM
Physalis_philadelphica_ASAT1     GMRANSSDRLLKPSILFQTMNLRPFIPLPEDAAGNFSSSLSVPTYVEEEM
Physalis_pruinosa_ASAT1          GMRANSSDRLLKPSILFQTMNLRPFIPLPEDAAGNFSSSLSVPTYVEEET
                                 ************************************************* 

Physalis_coztomatl_ASAT1         KLSRLISELRKGKEQLSDKYRKCKGNPQELVDTTMRSFQEIRDLFKDTDF
Physalis_philadelphica_ASAT1     KLSRLVSELRKGKEQLSDKYRKCKGNPQELVDTTMRSFQEIRDLFKDTDF
Physalis_pruinosa_ASAT1          KLSRLVSELRKGKEQLSDKYRKCKGNPQDLVDTTMRSFQEIRDLFKDTDF
                                 *****:**********************:*********************

Physalis_coztomatl_ASAT1         DLYRCSSLANYPLYDVDFGWGKPDNVISMVDYPLRNIFSLYDNKTCDHIE
Physalis_philadelphica_ASAT1     DLYRCSSLANYPLYDVDFGWGKPDNVISMVDYPLRNIFSLYDNKTCDHIE
Physalis_pruinosa_ASAT1          DLYRCSSLANYPLYDVDFGWGQPDKVMSMVDYPLRNIFSLYDNKTGDHIE
                                 *********************:**:*:****************** ****

Physalis_coztomatl_ASAT1         AQVSLDEESKMSALLREMGQLIQFKLVDE
Physalis_philadelphica_ASAT1     AQVSLDEESKMSALLREMEQLIQFKLVMN
Physalis_pruinosa_ASAT1          AQVSLDEENKMSAFLREMEQLIQFKLVDE
                                 ********.****:**** ******** :
```

**Supplementary Figure 15 – Full amino acid alignment of characterized *Physalis* ASAT1s.** Alignment was created using CLUSTALW. All three proteins are 429 amino acids in length. Catalytic motifs are labeled with a blue. Asterisks indicate complete conservation of a residue, with all sequences having the same amino acid. Colons indicate differing amino acids with strongly similar properties, periods indicate differing amino acids with somewhat similar properties, and spaces indicate differing amino acids that are not similar.

| retention (min) | %A (10mM ammonium formate) | %B (ACN) |
|---|---|---|
| 0 | 95 | 5 |
| 6 | 25 | 75 |
| 12 | 15 | 85 |
| 13 | 10 | 90 |
| 14 | 0 | 100 |
| 16 | 0 | 100 |
| 16.1 | 95 | 5 |
| 20 | 95 | 5 |

**Supplementary Figure 16 – LC-MS acylsugar gradient.** Gradient was used for acylsugar assays and acylsugar extracts, in addition to both positive and negative mode. Solvent A is 10mM ammonium formate adjusted to pH 2.8 with formic acid, solvent B is 100% acetonitrile (ACN). Flow rate was set at 0.3 mL/min. Additional parameters listed in methods.

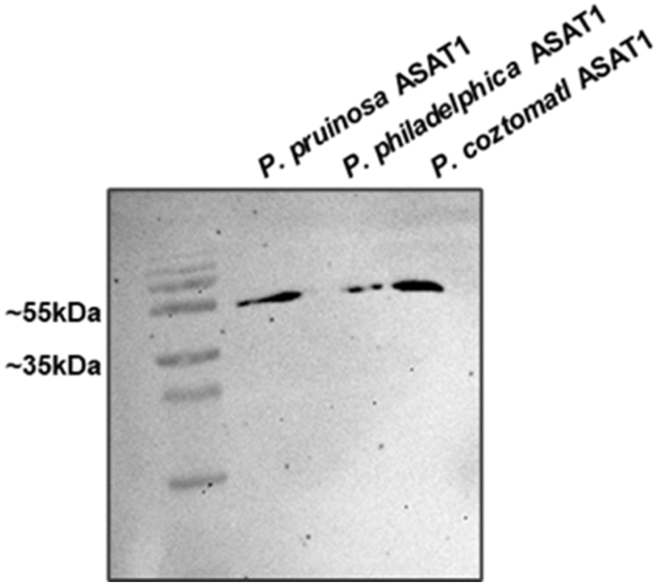


**Supplementary Figure 17 – Immunoblot of purified ASAT1s.** Immunoblot analysis was performed using a histidine tag antibody showing purification of *Physalis* ASAT1s
